# Culture-free generation of microbial genomes from human and marine microbiomes

**DOI:** 10.1101/263939

**Authors:** Alex Bishara, Eli L. Moss, Mikhail Kolmogorov, Alma Parada, Ziming Weng, Arend Sidow, Anne E. Dekas, Serafim Batzoglou, Ami S. Bhatt

## Abstract

Our understanding of natural microbial communities is shaped by the careful investigation of a relatively small number of isolated and cultured organisms, and by analysis of genomic sequences obtained by culture-free metagenomic sequencing approaches. Metagenomic shotgun sequencing has facilitated partial reconstruction of strain-level community structure and functional repertoire. Unfortunately, it remains difficult to cost-effectively produce high quality genome drafts for individual microbes without isolation and culture. Recent molecular techniques that partition long DNA fragments and then barcode short fragments derived from them produce “read clouds”, which are short-read sequences containing long-range information. Here, we present a novel application of a read cloud technique to microbiome samples, as well as Athena, a *de novo* assembler that uses these barcodes to produce improved metagenomic assemblies. We apply our approach to sequence human stool samples from two healthy individuals, and compare it to existing short read and synthetic long read metagenomic sequencing approaches. We find that read cloud metagenomic sequencing and Athena assembly produce the most complete individual genome drafts. These genome drafts are also highly contiguous (>200kb N50, <10 contigs), even for bacteria that have relatively low (20x) raw short read sequence coverage. We also apply this approach to a significantly more complex marine sediment sample and obtain 23 genome drafts with valuable 16S ribosomal RNA taxonomic marker sequences, nine of which are complete genome drafts. Read cloud metagenomic sequencing allows culture-free generation of high quality microbial genome drafts using only a single shotgun experiment.

## Introduction

Short-read sequencing and assembly have played an instrumental role in advancing the study of bacterial genomes beyond the minority of organisms that have been isolated and cultured^1^. This has greatly expanded our understanding of the genomic structure and dynamics of complex microbial communities that range from the human microbiome^2–4^ to environmental communities in the ocean, soil, and beyond^5–8^. However, the precise gene coding potential and consequent functional capabilities of organisms within these complex systems remains poorly understood.

Despite large-scale sequencing efforts, analysis of sequences from diverse environmental samples has revealed that major novel taxonomic lineages are entirely unrepresented in current reference collections^9, 10^, such as Refseq^11^. For example, even prevalent clades within heavily sequenced niches, such as Clostridiales and Bacteroides within the human gut, do not currently have a collection of isolate reference genomes that are fully representative of those identified with metagenomic shotgun sequencing^4^. Thus, there is a critical need for methods that will accelerate the generation of high quality genome drafts from shotgun sequencing of microbiome samples. Such genome drafts will greatly facilitate both the taxonomic and functional characterization effort of currently poorly understood clades, and also allow us to better understand genome variability within strains of a given species.

Specialized computational techniques have been developed to generate draft genomes for individual organisms within sequenced metagenomic samples. These techniques include dedicated metagenomic assemblers^12–14^ and metagenome draft binning based on sequence similarity^15–18^ and coverage depth covariance^19,20^. Binning techniques can group assembled sequences into significantly more comprehensive drafts, but these techniques cannot properly assign sequences that are shared between multiple bacterial strains. Sequencing reads produced by existing high throughput platforms (typically 100-250 base pairs) are too short to span many types of shared or duplicated sequences, and as a result, regions containing these types of sequences remain unassembled. Such shared and duplicated sequences arise from two main biological sources: (1) closely related organisms that underwent divergent evolution can share potentially many genome regions; (2) closely related or unrelated organisms can share sequences obtained by horizontal gene transfer and movement of mobile genetic elements^21,22^. These types of sequences pose challenges for short-read metagenomic sequence analysis, representing an inherent limitation of current short read sequencing approaches. To overcome this challenge, a complementary molecular approach to assemble these classes of genomic sequences is necessary.

In principle, long-read sequencing approaches can be used to address these issues. Long-read platforms such as Pacific Biosciences’ Single Molecule Real Time sequence approach have been successfully applied to close genomes of cultured isolates^23–25^ and dominant organisms within more complex mixtures^26^. However, these single molecule platforms have lower throughput and a higher error rate in comparison to short reads. These single molecule platforms also typically require higher input DNA mass (~100ng), which bars application of these platforms to biological samples where high molecular weight DNA is limited.

Synthetic long read (SLR) approaches such as Illumina Truseq Synthetic Long Reads^27^ have also been used to improve metagenomic assemblies^28,29^. The SLR approach produces long read sequences by partitioning long (~10kbp) fragments of DNA, clonally amplifying these fragments, and then shearing and generating a uniquely barcoded short-read sequencing library from each population of clonally amplified long fragments. Sequencing of the resulting library yields deep (50x) short-read coverage of the original long input DNA molecules, such that each can be assembled into a virtual long read. These virtual long reads are further assembled by overlap-layout-consensus assembly. Very deep sequencing applied to a healthy human stool sample using this SLR approach has allowed draft genome assembly for a subset of bacteria from that given sample^30^. Other applications of SLR sequencing to more complex environmental samples have relied on binning and direct examination of the virtual long read sequences to gain insight after encountering difficulty assembling them into longer genome sequences^29,31,32^. The lack of improvement in draft quality using SLR sequencing in studies of more complex environmental samples is most likely due to higher species richness and evenness in representation, which leads to individual genomes being covered with an inadequate number of virtual long reads. The challenges in overall throughput and high cost associated with the SLR approach have likely hindered its adoption and generalized utility.

The recent 10X Genomics platform streamlines the short-read barcoding process by utilizing more than a million droplet partitions to yield uniquely barcoded short-read fragments from one or a few long molecules trapped in each droplet partition^33^. Sequencing of libraries generated by this platform yields shallow-coverage groups of barcode-sharing reads, which we will refer to as read clouds^34^ (also referred to as linked-reads^33^). Though both read cloud and SLR approaches use long fragment partitioning, read clouds trade off shallower short-read coverage of each individual long fragment for an increase in the total number of long fragments sequenced (see Supplementary Discussion on read cloud and SLR sequencing). The 10X Genomics platform thus offers an attractive combination of long-range information, high throughput, high nucleotide accuracy, and low input mass requirements. This platform and similar ones predating it have demonstrated utility for this approach in reference-based human haplotype phasing^28,33,35–37^, and also in resolving complex structural variations in human genomes^38^. To date, their potential for *de novo* metagenomic sequence assembly has yet to be explored.

We present, to our knowledge, the first application of read clouds provided from the 10X Genomics platform to sequence human and marine microbiome samples. We introduce an assembler, Athena, that uses the barcode information from read clouds to produce high quality genome drafts from a single shotgun sequencing experiment. We first test our approach on a mock metagenome consisting of a staggered mixture of genomic DNA from 20 known bacterial strains, and use Athena to accurately assemble and place multiple copies of the ribosomal RNA (rRNA) operon within the draft assembly of each strain. We then apply our technique to sequence the gut microbiome of two healthy individuals, and compare our approach to existing short read and SLR approaches. We find that our approach combines the advantages of both short read and SLR approaches, and is capable of producing many highly contiguous drafts (>200kb N50, <10 contigs) with as little as 20x raw short-read coverage. Lastly, we apply our approach to a significantly more complex marine sediment sample and generate 23 genome drafts that contain valuable 16S taxonomic marker sequences, nine of which are also complete.

## Results

### Read cloud sequencing and Athena Assembly

We developed the Athena assembler to use long-range information encoded within barcoded short-read sequences. In our approach, we extract long DNA fragments and use the 10X Genomics Chromium platform to obtain barcoded short reads for our samples (Figure 1a). The resulting short reads are first stripped of their barcodes and jointly assembled using a standard short-read assembler (see Methods) to obtain an initial assembly of the metagenome in the form of sequence contigs. These seed contigs are then provided to the Athena assembler for further metagenome sequence assembly (Figure 1b). The same barcoded short reads are mapped back to the seed contigs and read pairs that span contigs are used to form edges in a scaffold graph. Branches in this scaffold graph correspond to ambiguities encountered by the short-read assembler. At each edge, Athena examines the short-read mappings together with the attached barcodes to propose a simpler subassembly problem of a pooled subset of barcoded reads that can potentially assemble through branches in the scaffold graph (see Supplementary Athena Description). The selection of this read subset removes the majority of reads considered during the initial assembly while retaining reads that cover the local target sequence, isolating the local subassembly problem from the broader metagenome. The much smaller and independent subassembly problems are performed separately for every edge in the scaffold graph to yield longer, overlapping subassembled contigs that resolve branches in the scaffold graph. The initial seed contigs and intermediary subassembled contigs are then passed as reads to the long read de Bruijn graph-based assembler, Flye^39,40^, which determines how to assemble the target genome from these much longer contigs. The resulting metagenome assembly consists of more complete sequence contigs resolving repeats that are too difficult to assemble with short-read techniques alone.

**Figure 1.**
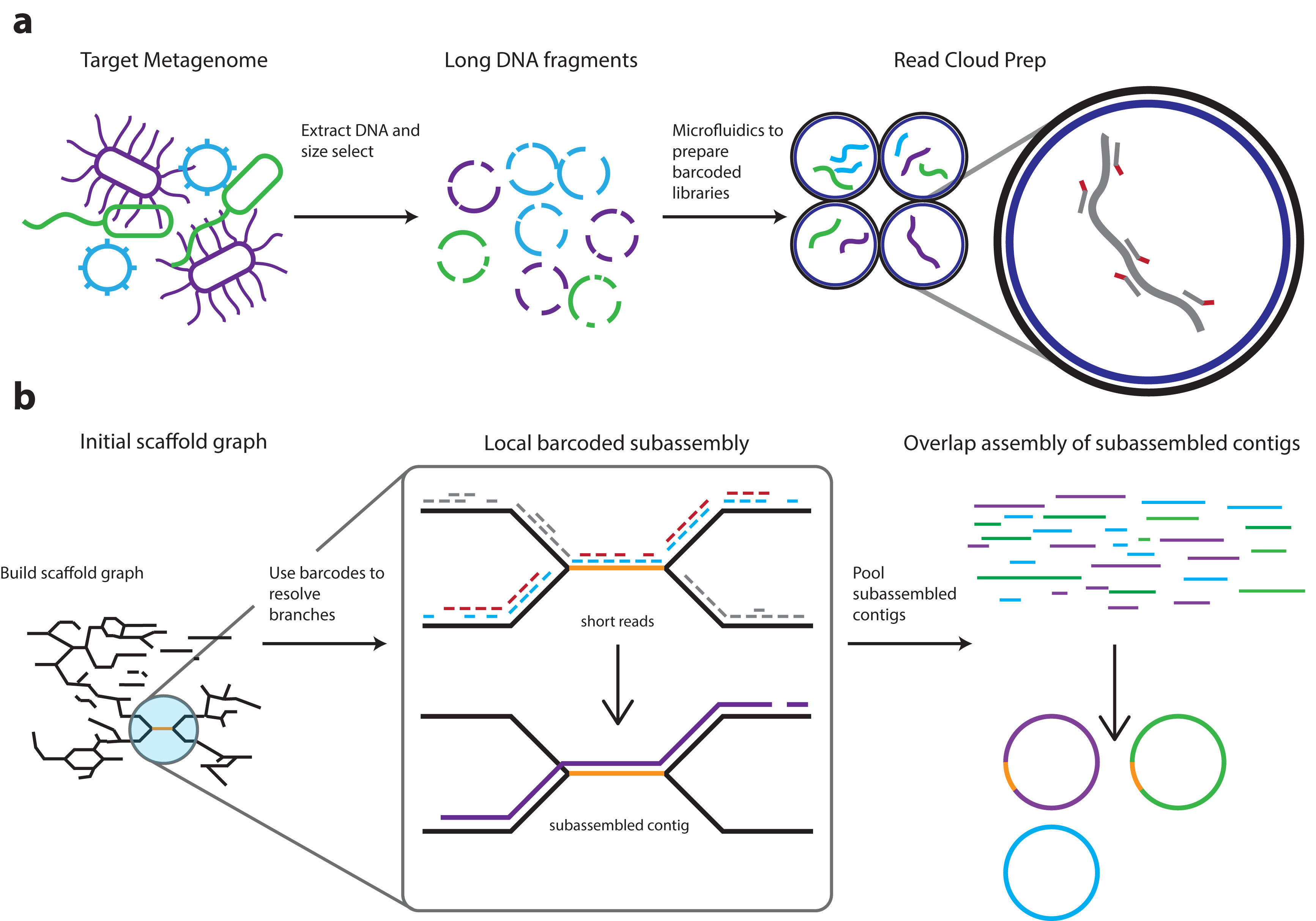
Overview of the read cloud shotgun sequencing and assembly approach. **a)** DNA is first extracted from microbiome samples and is size selected to enrich for long DNA fragments. The long fragments are then diluted and undergo sparse partitioning across more than a million droplet partitions (using, for example, the 10X Genomics Chromium library preparation platform). Degenerate amplification of these long fragments is then performed within these partitions to obtain barcoded traditional libraries -- each with a barcode unique to its partition. These libraries are then pooled and sequenced with an lllumina instrument. **b)** The Athena assembler uses read clouds to yield more complete drafts, in which genomic repeats are also accurately placed. An example repeat that is resolved and placed by Athena is shown in orange. 1) Read clouds are first assembled with standard short-read techniques (e.g. metaSPAdes, IDBA-UD, MEGAHIT) to obtain seed contigs, input reads are mapped back to these seed contigs, and read pairs that span two seed contigs are used to build a scaffold graph containing unresolvable branches. 2) At each edge, Athena proposes a much simpler subassembly problem on a pooled subset of barcoded reads informed by the scaffold graph mappings. Example short reads with red and blue barcodes are passed to a short-read assembler to perform subassembly, which yields a longer subassembled contig that disambiguates branches in the scaffold graph. 3) The resulting subassembled contigs, together with the initial seed contigs, are then passed to as reads to the long read de Bruijn graph based assembler Flye for final assembly. The resulting draft assembly metagenome produces more complete and more contiguous drafts in which repeats are also assembled and correctly placed.

### Assembly of a mock metagenome community

As a first validation of our approach, we applied Athena to assemble a read cloud library of a staggered mixture of genomic DNA from 20 bacterial strains (ATCC MSA-1003, see Methods), 19 of which have closed reference genomes available that allow us to assess the accuracy of our approach. There is currently no closed reference genome available for the *Pseudomonas aeruginosa* strain present in this mixture. The read cloud library was sequenced on one full lane of an Illumina HiSeq 4000 sequencer yielding roughly 74Gb of raw short-read sequences. We assessed the Athena assembly against the available references and also evaluated Athena’s ability to accurately assemble the conserved 16S and 23S genes of the ribosomal RNA operon (which is 5-7kb in size depending on the size of the spacer between 16S and 23S genes). These genes are also highly conserved across all bacterial species, yet contain sufficient divergence between species, allowing for their use as an informative marker for phylogenetic characterization of microbial communities^41^. These interspecies repeat units, which also occur in multiple nearly identical copies within each individual genome^42,43^, serve as a useful model of the assembly issues created by duplicated and conserved sequences.

We assembled the read cloud library of the 20 strain mixture using Athena and evaluated the overall draft quality against the known reference genomes. In order to compare against conventional short-read assembly, we also assembled the raw barcode-stripped read cloud sequencing data using a standard short-read assembler. The assembled metagenome drafts of each approach were evaluated using MetaQUAST^44^ to assess contiguity, base-error rates, and mis-assemblies (Supplementary Table 1). MetaQUAST produces these performance statistics by aligning genome drafts against available closed reference genomes to discover sequence differences with respect to a given reference. Athena-assembled drafts were significantly more contiguous than short-read assembled drafts with a median contig N50 increase of 7.6-fold for organisms with a minimum of 20x raw short read coverage. This contiguity was achieved without sacrificing overall accuracy when compared against conventional short-read assembly. We found Athena assembly to be comparable to short-read assembly on two important metrics: base-error rates (8.97 vs. 10.45 mismatches per 100kbp) and also the total number of mis-assemblies (67 vs. 61). Two draft genomes were excluded from mis-assembly analysis: *P. aeruginosa*, as there is no available closed reference genome for the strain sequenced in the mock metagenome; and *Acinetobacter baumannii*, as the available reference genome was determined to have several mis-assemblies following subsequent rRNA operon analysis.

We then identified 16S/23S rRNA operons within drafts from both approaches, and compared the placement of these repeats against the known reference genomes to ensure correct placement. We used RNAmmer^45^ to find completely assembled rRNA operons with at least 3kbp of flanking sequence within the short read and Athena drafts. Conventional short-read assembly was unable to correctly assemble and place a single rRNA operon. In contrast, Athena read cloud assembly produced 41 copies of the complete rRNA operon across multiple species (Supplementary Table 2). Three of the four copies from *A. baumannii*, two of the three from *Propionibacterium acnes*, and one of the four from *Rhodobacter sphaeroides* were determined to have differing placements in their respective genome drafts when compared to the closed reference genomes. For *A. baumannii* and *P. acnes*, inspection of the raw read cloud alignments against the reference genomes suggested the possibility that these closed reference genomes may also be mis-assembled. Long-range PCR and Sanger sequencing confirmed that the three discordant *A. baumannii* rRNA operons were misassembled in the available closed reference genome (Genbank ID: CP000521.1) and correctly assembled using read clouds (Supplementary Figure 1). Long-range PCR and Sanger sequencing confirmed only one of the three assembled rRNA operons and PCR did not yield amplification of the other two (Genbank ID: CP003084.1, Supplementary Figure 1). Further inspection of read cloud alignments to the *P. acnes* reference genome suggests the presence of a fourth rRNA operon in tandem with one of the other three in this organism, which would complicate correct assembly by any approach. All 41 assembled rRNA operons were correctly assigned to their respective genome and only three, two from *P. acnes* and an additional one from *R. sphaeroides*, were determined to be misassembled.

Athena read cloud assembly facilitated the assembly and proper placement of the 16S/23S rRNA operon for the organisms sequenced in the mock metagenome and yielded significant improvements in overall draft quality.

### Assembly of genomes from the human intestinal microbiome

To test the generalizability of this approach to natural biological samples, we next applied read cloud sequencing and Athena assembly to stool samples from two healthy human participants, P1 and P2. To assess the effects of different DNA extraction methods on long fragment assembly performance, we used Puregene enzymatic cell lysis to extract DNA from sample P1 and Qiagen mechanical cell lysis to extract DNA from sample P2. In order to properly evaluate performance against alternative metagenomic sequence assembly approaches, we also prepared standard Illumina Truseq short read and Illumina Truseq SLR sequencing libraries from extracted DNA. Read cloud and SLR library preparations both require long DNA fragments whereas Truseq library preparation does not. Thus, extracted DNA to be used in read cloud and SLR libraries was first subjected to size selection (see Methods, Supplementary Table 3). For each stool sample, prepared short read Truseq and read cloud libraries were multiplexed together and sequenced using an Illumina HiSeq 4000 sequencer yielding roughly 40Gb of raw short-read sequences per library. SLR libraries cannot be multiplexed, so instead each of the two SLR libraries was given its own full lane of sequencing on a HiSeq 4000 yielding roughly 102Gb of raw short-read sequences for each library (Supplementary Table 4).

Genus-level community compositions for each of the three sequencing approaches were first assessed using k-mer based short-read classifications (Figure 2a,b and Supplementary File 1). The three sequencing approaches yielded comparable community compositions within each stool sample. We further performed short-read classification of additional libraries from several low-pass sequencing experiments to test whether the method of DNA extraction (enzymatic vs. mechanical bead beating) or size selection had any major effect on genus-level community composition. Though some less abundant genera differed in their abundance rank, the community composition was largely concordant between all approaches tested (Supplementary Figure 2).

**Figure 2.**
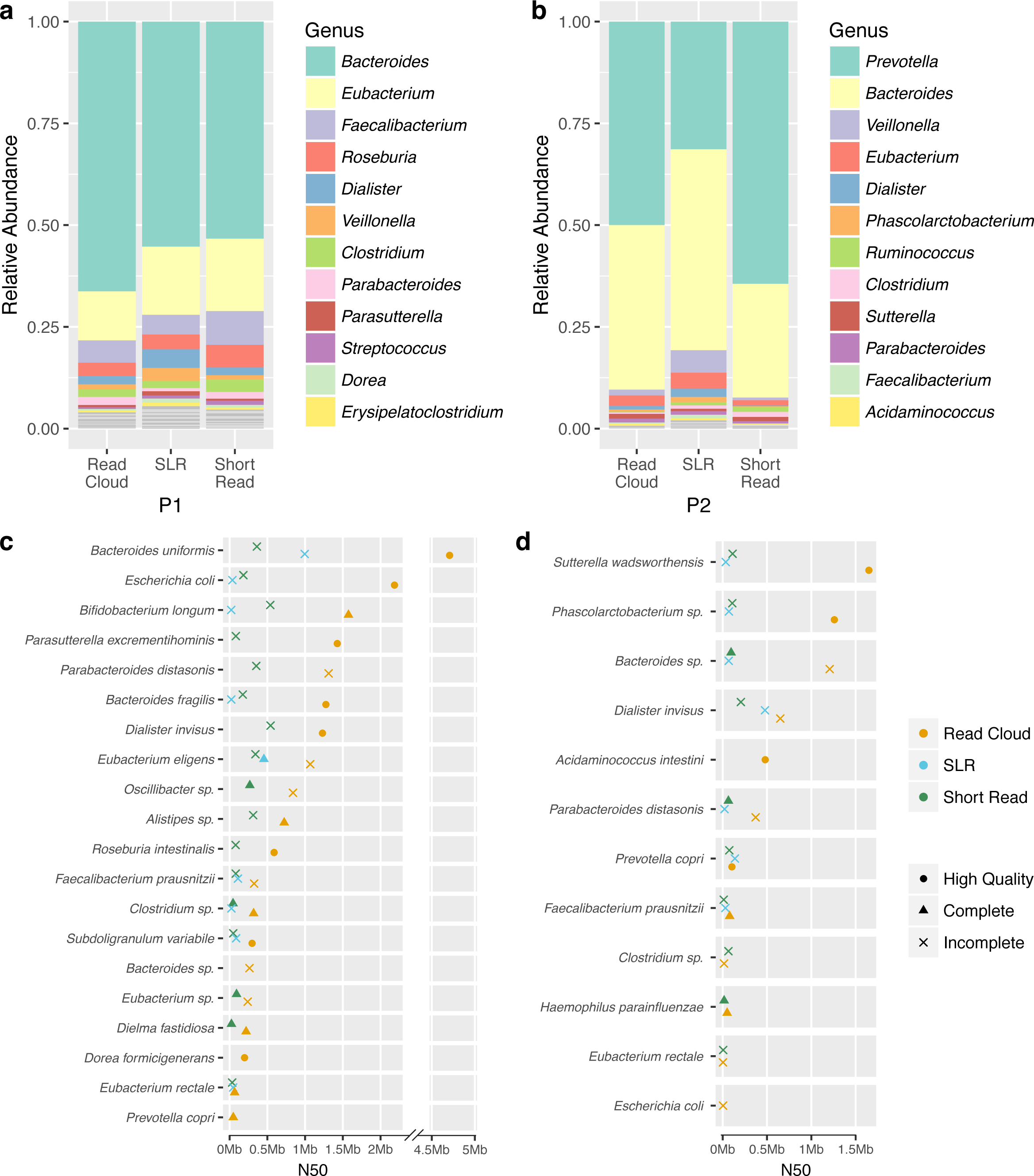
Composition of stool microbiome communities from two healthy human participants. a, b) Relative abundances of genera as determined by short-read classification for each of the three libraries from samples P1 and P2. The relative representation of genera appears is fairly concordant between the three different library preparation methods (short read, SLR, read cloud) for each sample. Sample P1 is more diverse than sample P2 at the genus-level. c, d) Comparisons of genome draft contiguity, as measured by N50, for taxa that were present in samples P1 and P2. The read cloud approach results in a larger number of more contiguous genome drafts than the short read or SLR approaches. Results are only displayed for the largest bin of each taxon determined to be present. The completeness and contamination of genome drafts for these taxa was determined by assessing the presence of lineage-specific single copy core genes as predicted by checkM. Genome drafts were designated as incomplete (‘x’, <90% completeness), complete (circle, >90% completeness and <5% contamination), high quality (triangle, complete and with at least 18 tRNAs, as well as at least one of each of the 5S, 16S, and 23S rRNA genes). Read cloud sequencing and assembly produces many high quality and complete drafts. The read clouse sequencing and assembly-produced drafts are much more contiguous as compared to those obtained from SLR and short read sequencing.

To compare performance of the three sequencing approaches, the appropriate assembly approach was applied to each sequenced library to obtain initial metagenomic drafts. Short read libraries were assembled using a conventional short-read assembler (see Methods) and the read cloud libraries were assembled using Athena. The SLR libraries required a two stage assembly process (see Methods), in which the barcoded short-reads are first assembled into virtual long reads using a modified short-read assembler. These virtual long reads are then assembled using overlap-layout-consensus assembly to yield a draft of the entire metagenome. Although SLR libraries received a high amount of total raw short-read sequence (102Gb for both P1 and P2), the total sequence in the form of virtual long reads was significantly lower than the amount of raw short-read sequence. (0.64Gb for P1 and 0.55Gb for P2, Supplementary Table 4).

Read cloud sequencing and assembly resulted in much larger microbial sequence contigs compared to both SLR and short-read sequencing and assembly. Nearly 144Mb of sequence from P1 and 40Mb of sequence from P2 were assembled using read clouds into contigs with a minimum size of 100kbp, compared to just 68Mb and 22Mb using short reads, and 26Mb and 14Mb using SLRs (combined results in Supplementary Figure 3). The overall size of the read cloud metagenome drafts was also significantly larger compared to the SLR metagenome drafts (345Mb vs 55Mb in P1 and 229Mb vs 31Mb in P2), highlighting the benefit of increased throughput of our approach that allows assembly of lower-abundance organisms.

To assess the ability of each approach to produce usable genome drafts for constituent bacteria, we binned metagenome draft contigs and used annotations of contigs to obtain genus-level and/or species-level assignments for each resulting bin (see Methods, Supplementary Figure 4, Supplementary Table 8). The resulting bins were assessed as genome drafts by the presence of lineage-specific single copy core genes to determine completeness and contamination. Using a previously described set of criteria, we refer to a genome bin as a complete genome draft if it is >90% complete and <5% contaminated as predicted by checkM^46^. We refer to the subset of these complete genome drafts as *high quality*, adopting a previously defined standard^47^, if the draft also contains at least 18 tRNA loci and at least one copy each of 5S, 16S and 23S.

Read cloud sequencing yielded complete and high quality genome drafts for bacteria from both samples P1 and P2 (Figure 2c,d). Our most contiguous, high-quality read cloud draft was for *Bacteroides uniformis* in sample P1, which was contained completely in three contigs of sizes 4.7Mb, 369kb, and 25kb. Several other bacteria from P1 were also well-assembled including *Bifidobacterium longum, Escherichia coli,* and *Bacteroides fragilis*. Read cloud sequencing on sample P2 yielded much fewer complete or high quality genome drafts, due to the reduced number of well-covered bacteria. Alignments of input short reads from the P1 and P2 read cloud libraries to genome bins allowed estimation of abundances of individual organisms, and confirmed sample P2 contained fewer and much more abundant bacteria (Supplementary Table 5). Read clouds generated genome drafts annotated to representative species, which belonged to genera determined to be abundant by direct classification of the raw short reads. In sample P1, read clouds also generated four complete drafts and one high quality draft annotated as belonging to the genus *Clostridium*, though none received species-level annotations. In addition, genome binning of the metagenome drafts also yielded multiple other near complete genome bins annotated as *F. prausnitzii*, but each of these were determined to have contamination as assessed by single copy core gene presence. These results suggest the presence of multiple bacterial genomes within sample P1 that are most closely related to the known species *F. prausnitzii*, yet each of these are distant enough from one another to be separately assembled into near complete drafts.

Though read cloud assembly and binning yielded a single high quality genome draft that was annotated as *Prevotella copri* in sample P2, the N50 of 103kb for this read cloud draft was notably low despite this bin having 2,836x short read coverage. Analysis of short reads originating from this genome bin in the read cloud library illuminated the unusual presence of five high-copy (>10 copies) genomic elements that likely impeded improvements in assembly by our approach (see Supplementary Methods, Supplementary Figure 5). These five high-copy genomic elements were annotated at either the nucleotide or amino acid level as putative transposases (Supplementary Table 6). Based on the copy number, these genomic elements are predicted to occur at many locations within the *P. copri* genome present in sample P2. One of these five transposase repeat elements, which had high sequence homology to known insertion sequence IS932, occurred at an estimated copy number of 63 within the *P. copri* genome present in sample P2.

We assessed the impact of different DNA extraction methods used on each stool sample on the resulting long DNA fragment size. The sizes of long fragments used as input for read cloud sequencing were estimated from the read cloud sequencing data (see Supplementary Methods). Fragments from sample P2, which was subjected to mechanical DNA extraction, had a median size of 5kbp. Fragments from sample P1, which was subjected to enzymatic DNA extraction, had a median size of 8kbp and were noticeably longer than those from sample P2 (Supplementary Figure 6). However, assembly of each read cloud library still yielded highly contiguous genome drafts such that the use of shorter fragments did not seem to significantly impact assembly quality.

The read cloud approach was superior to both the short read and SLR approaches in its ability to recover genome drafts for individual bacteria (Figure 3). The combined results from read cloud sequencing of samples P1 and P2 yielded a total of 27 complete drafts, compared to 18 from short read sequencing. SLR sequencing produced only one complete draft despite receiving twice the amount of raw short read sequencing for each sample (due to the inability to multiplex libraries). Read clouds produced the most complete drafts that were also highly contiguous (N50 > 200kb) with a total of 16, compared to just one each from short read and SLR approaches. For each approach, raw input short reads were mapped back to their respective sequence contigs in order to estimate the total short read coverage used for the assembly of each organism. Read clouds were able to produce complete genome drafts, a large fraction of which were also highly contiguous, with as little as 20x short read coverage for some bacteria (Figure 3b, 3c). The short read approach also produced multiple complete drafts at low coverage. However, the resulting drafts from short reads were fragmentary compared to the read cloud drafts, even for bacteria with high short read coverage. Though short reads can produce complete genomes, they are incapable of producing high quality genomes as short fragment length limitations prevent assembly of 16S/23S rRNA operons. The SLR approach produced only a single draft that was complete and highly contiguous (N50 > 200kb). This well-assembled organism in the SLR library happened to have over 2,300x short read coverage, and the SLR approach did not produce any other well-assembled genome drafts for organisms with less short read coverage. Of all three tested approaches, read clouds were the only approach capable of producing complete drafts that were also high quality, in which at least 18 tRNA loci and at least one copy each of 5S, 16S and 23S were assembled (Figure 3d,3e,3f).

**Figure 3.**
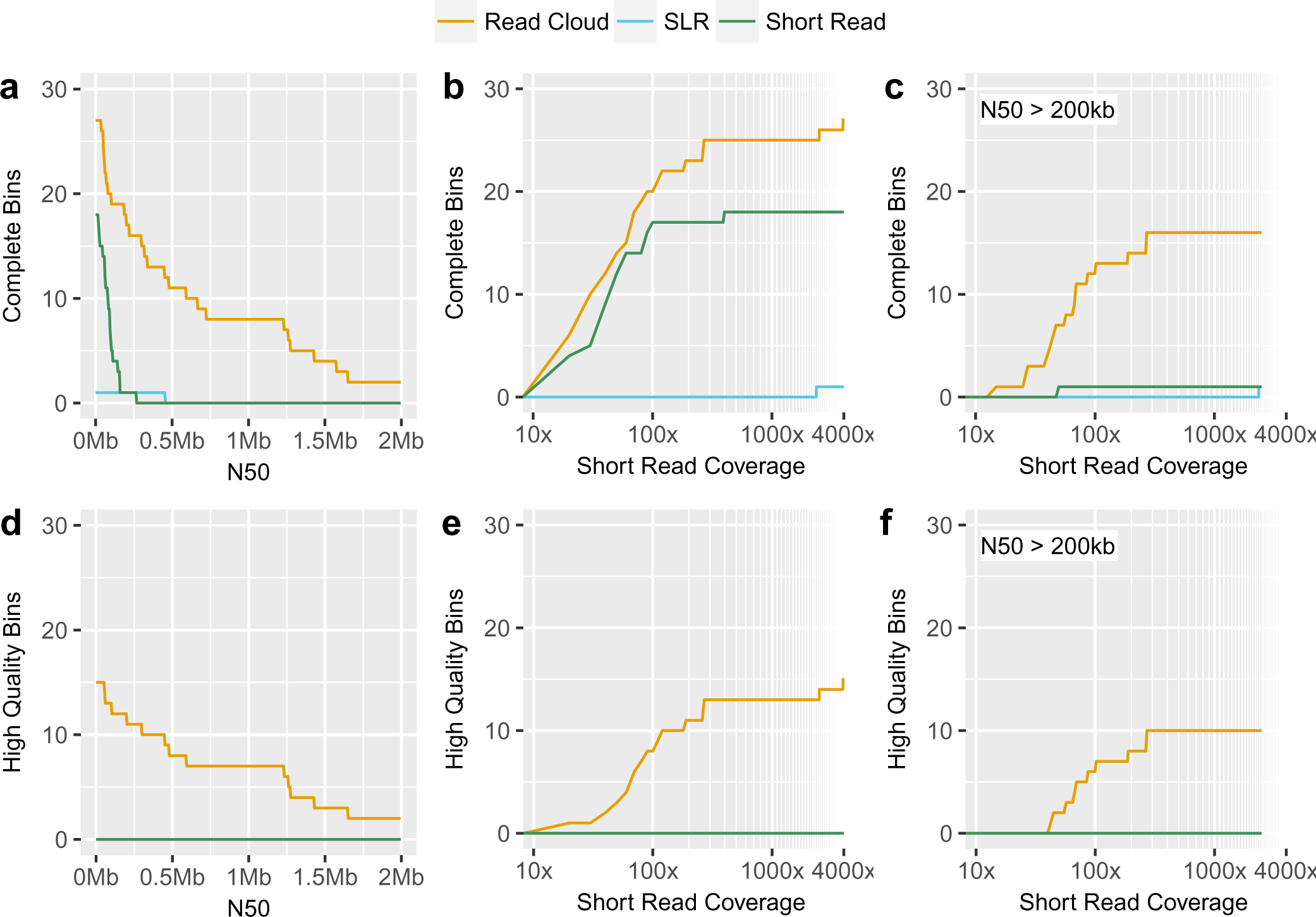
Combined genome draft results of read cloud, SLR, and short read approaches applied to healthy human stool samples. Under various performance metrics, read clouds (gold) consistently display superior performance in their ability to produce many complete and high quality genome drafts as compared to either SLRs (blue) or short reads (green) approaches. Performance was also superior even in low short read coverage regimes (defined as <50x coverage). Counts include all complete/high quality genome bins for all taxa in each approach. a) Number of complete genome bins (>90% completeness, 5% contamination) with a minimum N50. b) Number of complete genome bins with a minimum short read coverage depth. Genome bins with lower short read coverage correspond to less abundant organisms. c) Number of complete genome bins with an N50 of >200kb and a minimum short read coverage depth. d) Number of high quality genome bins (complete and with at least 18 tRNAs, as well as at least one instance each of the 5S, 16S, and 23S rRNA genes) with a minimum N50. e) Number of high quality genome bins with a minimum short read coverage depth. f) Number of complete genome bins with an N50 of >200kb and a minimum short read coverage depth.

We next assessed differences between the three approaches in their ability to produce complete drafts for particular taxa (Figure 4). Read clouds produced by far the most complete and high quality genome drafts in which all contigs were clustered into a single bin. In contrast, contigs from short reads for these single genome taxa were most frequently split across two or more bins. The only taxon for which SLRs produced a complete draft within a single bin, *Eubacterium eligens*, was not placed in a single bin using either read clouds or short reads. We identified all open read frames in each metagenome draft and used these annotations to evaluate the total gene content of each genome bin (see Methods). In one case, the SLR genome bin corresponding to *B. uniformis* had more genes than the corresponding read cloud draft. However, this bin was determined to be 15% contaminated and is likely to incorrectly contain genes from other organisms. For the majority of taxa discovered in samples P1 and P2, read clouds successfully binned more genes together than either short reads or SLRs.

**Figure 4.**
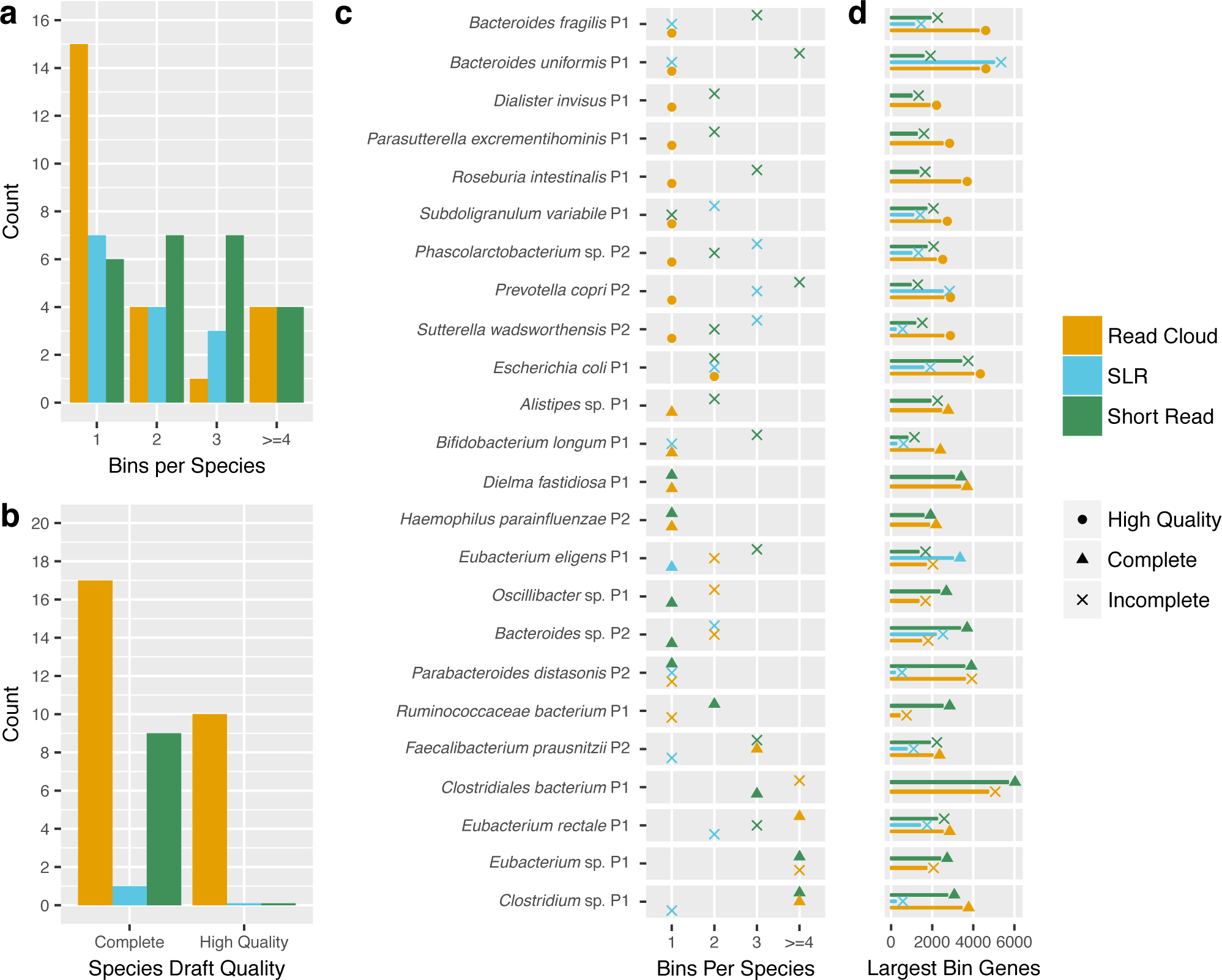
Completeness of genome bins produced by read cloud, SLR, and short read sequencing for various taxa present in healthy human stool samples. Read clouds (gold) consistently yield more complete and high quality genome drafts for taxa within singleton bins, as compared to SLR (blue) and short read sequencing (green), both of which split sequence contigs from single genomes into two or more genome bins. Taxa are only shown if it is represented in at least two approaches and at least one approach produced a complete bin. a) Counts of the number of bins containing sequence for each taxon for each of the three approaches. Read clouds produced the most singleton bins for the taxa considered. b) Counts of complete and high quality drafts for each approach. Read clouds produced the most complete genome drafts in singleton bins with 14. Ten of the 14 singleton bin complete genome drafts were designated as high quality. c) For each approach, the total number of genome bins annotated as belonging to a particular taxon. The largest bin produced by an approach for a particular taxon is designated as a incomplete (‘x’), complete (circle), or high quality (triangle) genome draft. For nearly all taxa that received a complete or high quality genome draft from a particular approach, only a single genome bin was annotated as belonging to these taxa. However, for some taxa, such as *Escherichia coli* and *Clostridiales bacterium*, these complete or high quality genome drafts were accompanied by a few much smaller incomplete bins that were also annotated as belonging to these taxa. d) Counts of the number of genes present in the largest bin for a particular taxon and approach. The read cloud approach yields the bins containing the largest number of genes for the majority of taxa. The SLR bin annotated as *Bacteroides uniformis* in sample P1 contains more genes, but was determined to be 15% contaminated. This suggests that such some of these genes assigned to the SLR bin for *Bacteroides uniformis* are likely from other organisms.

Comparisons of our high quality drafts against available closed reference genomes show both cases where genome structure is largely maintained, and also cases where large structural rearrangements are apparent (Figure 5). Both *Dialister invisus* and *E. eligens* were present and assembled into high quality genome drafts in both samples P1 and P2. Alignments of both *D. invisus* drafts from samples P1 and P2 illustrated large scale rearrangement with respect to the available reference genome. Inspection of these reference alignments indicates that the *D. invisus* strains generated by the read clouds in each sample are largely structurally divergent from each other as well. Interestingly, the draft recovered for *E. eligens* from sample P2 was structurally similar to the reference genome, whereas the draft recovered from sample P1 displayed two large scale inversions. Despite structural concordance in most our assembled drafts to the available reference genomes, all of them deviated substantially from the available references in sequence identity for alignable bases and also the total number of bases that were unalignable (Supplementary Table 7). The median nucleotide sequence identity was 98.5% and the median fraction of reference-unaligned bases in each draft was 15.7%.

**Figure 5.**
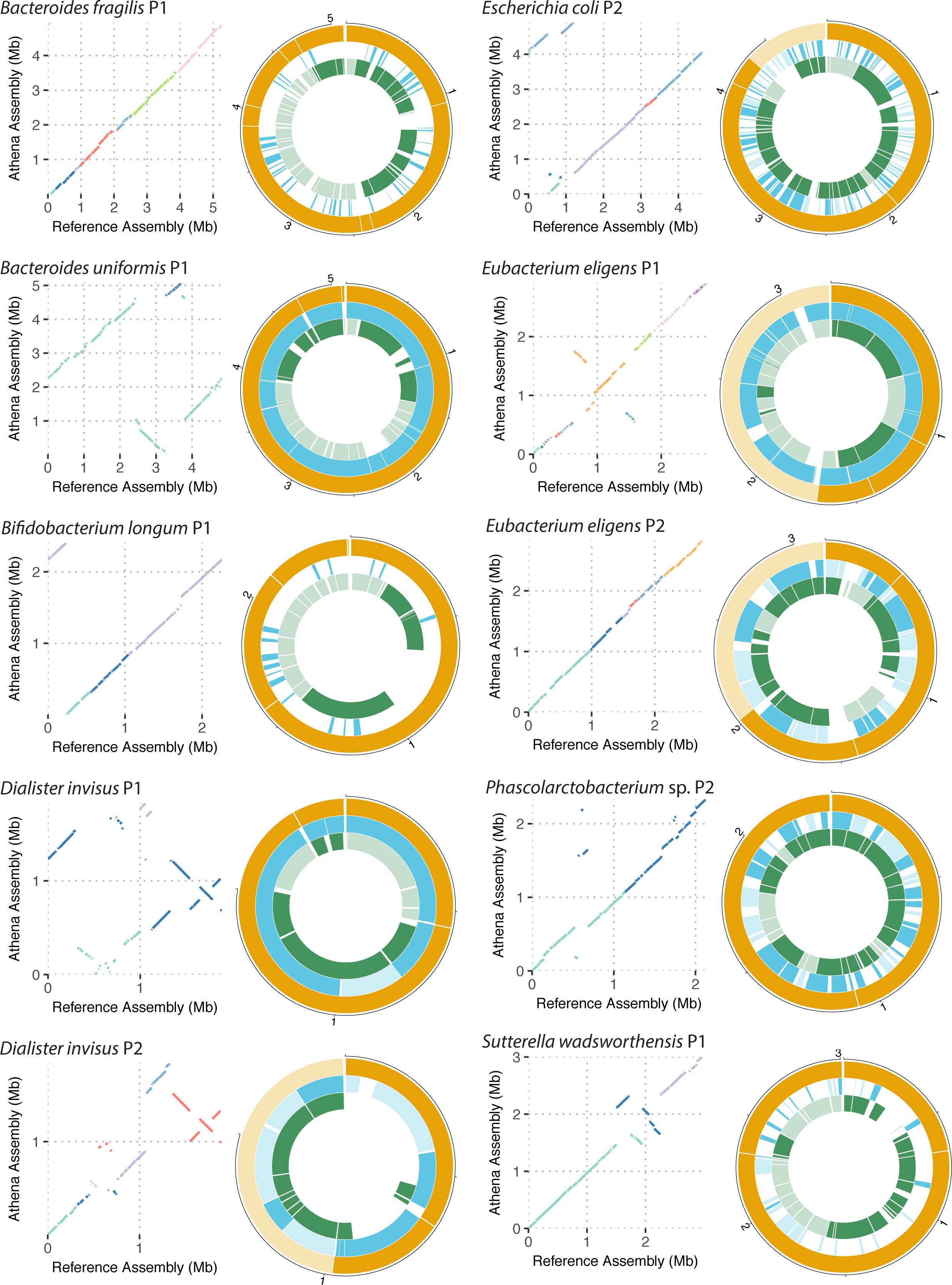
Comparisons often representative read cloud genome drafts to available reference genomes, and corresponding short read and SLR drafts. Dot-plot alignments between read cloud drafts (*y*-axis) and the closest available reference genome (*x*-axis) are shown. For each dot-plot, a given color corresponds to the alignment of a single contig in the read cloud draft against the available reference. Large-scale structural concordance and also differences including inversions are visually apparent. Alignments of SLR and short read drafts to the read cloud drafts for each taxon are also shown. In all cases, read cloud drafts were the most contiguous. For each approach, contigs belonging to the largest genome bin for a particular taxa are given a darker color, and the rest of the contigs in other bins are represented with a lighter color.

For the organisms assembled into high quality drafts using read clouds, alignments of the corresponding SLR and short read drafts illustrate the fragmentary nature of the drafts recovered by these two approaches. Organisms that were not present at high enough abundances within each of the samples received only sparse virtual long read coverage in the SLR libraries, such that further sequence assembly of these virtual long reads into a sequence contigs was generally not possible. Although the short read approach did not suffer from the same throughput limitation, it was nonetheless only capable of producing fragmentary genome drafts. The read cloud approach was the only one capable of producing high quality and highly contiguous genome drafts de novo from the studied human stool samples.

### Assembly of a marine sediment microbial community

To test the ability of read clouds to generate genome drafts from samples that are significantly more complex than human stool microbiomes, we applied read cloud sequencing and Athena to a marine microbiome. A deep-sea sediment core was extracted from superficial sediment at a water depth of 3535 meters using an MC-800 multi-corer. The sample was collected during an expedition in the Pacific Ocean approximately 115 kilometers off the northern California coast near San Francisco. DNA was extracted from this sample using a combination of mechanical bead-beating based and chemical lysis, and subjected to a size selection to enrich for long DNA fragments (see Methods). A read cloud library was prepared and sequenced on one full lane and a quarter lane on an Illumina HiSeq 4000 flow cell, yielding roughly 72Gb of raw short-read sequences (Supplementary Table 4). In order to successfully assemble this sample, which is significantly more complex than our human stool samples, we applied a specialized short-read assembler designed for use with large and complex metagenomes (see Methods). Modifications were also made to Athena to successfully assemble the sequencing data using the read cloud barcode information (see Methods).

The short-read assembled metagenome was 5.3Gb, indicating a very high species-richness in our marine microbiome. However, mapping of the input short reads back to this metagenome draft allowed us to determine that most of the assembled sequences belong to a large number of rare organisms each present at less than 0.1% relative abundance. Of the 5.3Gb that were assembled, only 517Mb of this sequence were present in contigs of at least 400 base pairs with short read coverage of at least 20x.

Athena read cloud assembly produced far more large sequence contigs and also many more assembled 16S ribosomal RNA (16S rRNA) taxonomic marker sequences than short-read assembly alone. Nearly 351Mb of sequence was assembled using Athena into contigs with a minimum size of 10kbp, compared to just 135Mb assembled using standard short-read assembly alone (Supplementary Figure 7). We next searched for assembled 16S rRNA loci within the short read and Athena metagenome drafts. Athena assembly yielded a total of 130 16S rRNA sequences compared to just 23 from short-read assembly.

We next assessed the ability of each assembly approach to produce usable genome drafts from the marine microbiome. Metagenome draft contigs were binned and each resulting bin was assessed as a genome draft by the presence of lineage-specific single copy core genes as with the human stool samples (see Methods, Supplementary Figure 4, Supplementary Table 9). Genome drafts were unable to be annotated with species-level or genus-level designations due to the current lack of representative isolate genomes for these microbiome samples.

Read cloud sequencing and Athena assembly consistently produced more genome drafts than short-read assembly alone (Figure 6). Athena assembly produced nine complete genome drafts, eight of which were also high quality. Short read assembly was unable to produce a single draft meeting these quality criteria. We next assessed the ability of each approach to produce intermediate quality genome drafts, which we defined as having >70% of predicted single copy core genes present and <10% of these genes present more than once. Athena produced 49 intermediate quality genome drafts of which 24 also contained assembled 16S rRNA sequences. Although short-read assembly produced 28 quality genome drafts, only six of these contained 16S rRNA sequences.

**Figure 6.**
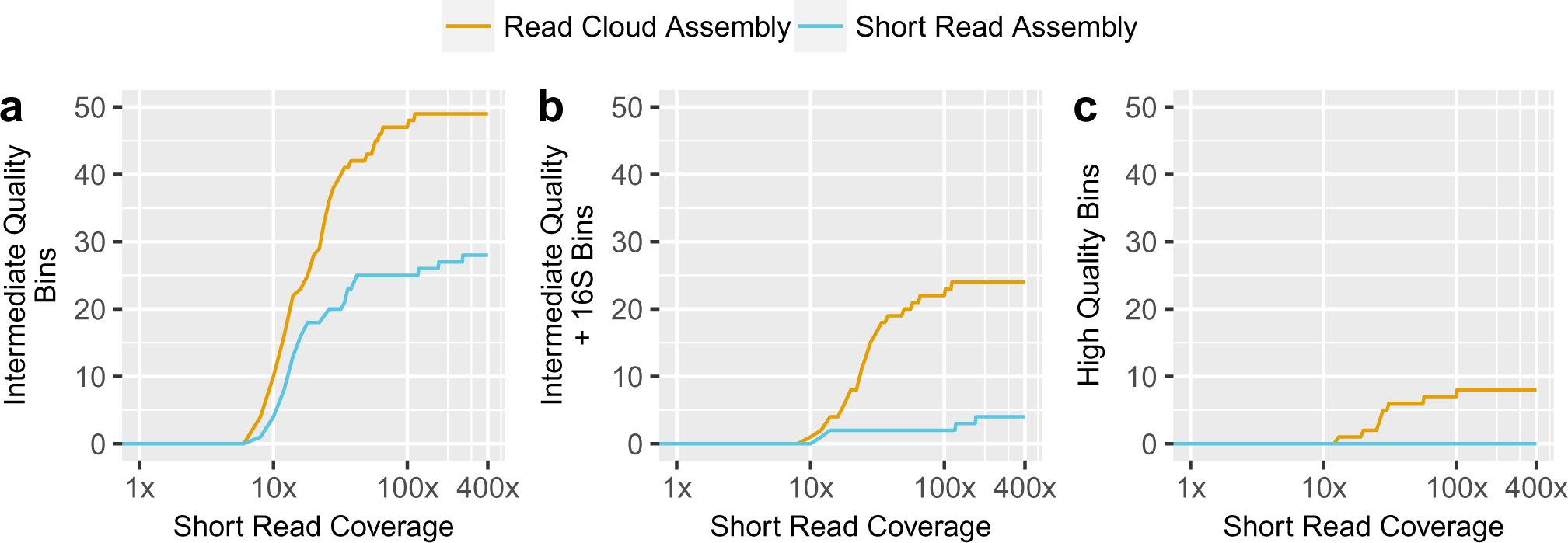
Genome draft results of Athena assembly and standard short-read assembly applied to read cloud sequencing of a marine microbiome. Athena read cloud assembly (gold) consistently produced the more genome drafts than standard short-read assembly (blue) with genome bins assessed as genome drafts under various different quality criteria. Athena read cloud assembly allowed significantly more 16S rRNA (16S) taxonomic sequences to be assigned to genome drafts than short-read assembly. The number of a) intermediate quality (>70% completeness and <10% contamination) genome drafts b) intermediate quality genome drafts with assembled 16S rRNA sequences, and c) high quality genome drafts with assembled 16S rRNA sequences with a minimum short read coverage depth are shown.

## Discussion

We present a novel approach using read clouds to generate high quality *de novo* genome drafts from microbiome samples with a single shotgun sequencing experiment. We demonstrate successful application of our read cloud approach with Athena assembly to generate high quality genome drafts within a mock metagenome sample, human stool samples from two healthy individuals, and a marine sediment sample. We compare our approach to alternative short read and SLR metagenomic sequencing approaches, and demonstrate that our approach can cost-effectively generate high quality genome drafts from complex microbiomes.

We anticipate that the approach presented here will benefit from future improvements in both DNA extraction techniques and long fragment barcoding approaches. Though our experiments show large scale concordance between overall community composition for long DNA fragments extracted under varying conditions, further improvements can likely be made to increase taxonomic concordance. Furthermore, relatively high input DNA mass is required when using existing mechanical cell lysis techniques, as a subsequent size selection to enrich for high molecular weight DNA incurs significant loss. Improved lysis techniques that preserve high molecular weight DNA from all present bacteria may reduce the effect of these techniques on community representation.

Read cloud sequencing and assembly helped shed light on prevalent gut commensals including those from the genera *Prevotella* and *Clostridia*. Several Clostridium strains from a single individual (P1) were assembled into complete drafts, suggesting that this approach will be valuable in characterizing strain populations of closely related taxa in human stool microbiomes. Our read cloud approach produced highly contiguous and complete genome drafts for nearly all bacteria with high sequence coverage present in our human stool microbiome samples. However, the complete genome draft produced for the most abundant organism in either sample, *Prevotella copri*, was more fragmented than might be expected given observed performance on less deeply covered organisms. This organism was determined to have several high-copy genomic repeat elements that likely complicated correct resolution of local genomic structure during subassembly in Athena. The Chromium platform currently groups several (~10) long fragments per barcode. Improvements that allow only a single long fragment per partition would greatly reduce the complexity of each subassembly task within Athena, and potentially allow read clouds to better assemble organisms with these high-copy repeats.

Further development of binning methods that take advantage of the read cloud barcode information will allow recovery of even more individual microbial genome drafts from the communities presented. Our current approach to produce individual genome drafts leveraged both our Athena assembler to significantly improve metagenomic contig assembly, as well as existing binning tools that were designed for use with conventional short read assembly techniques. These binning tools cluster contigs into groups with use of sequence metrics based on features such as nucleotide composition similarity (tetramer frequencies) and coverage depth. Although application of these tools worked well when applied to our improved metagenome draft contigs, they were unable to properly deconvolve a few members of some genera in our stool microbiome samples -- such as *Bacteroides* and *Faecalibacterium* -- and likely members of many less characterized genera within our marine sediment samples. Multiple genomes belonging to each of these taxa are likely present in similar abundances and have similar nucleotide compositions, such that the current metrics do not allow contigs from these taxa to be correctly separated into individual draft genomes. Read clouds that are mapped back to an assembled metagenome draft can provide linkage information for pairs of sequence contigs sharing barcodes. Sequences sharing many barcodes are indicative of sequences originating from the same input DNA fragments, which should then be binned together. Binning approaches that aim to incorporate of this linkage information will likely provide a stronger signal that can further disentangle closely related taxa within complex metagenomic samples.

Of the studied methods, our read cloud approach was the only one capable of generating genome drafts for the significantly more complex marine sediment sample. Our use of existing binning approaches yielded 42 partial drafts, of which 23 were tagged with informative 16S taxonomic marker sequences, and nine of which were determined to be complete drafts. Extensions of these binning approaches to use the linkage information present in the read clouds will likely allow the generation of far more complete bins from these complex samples. Further applications of our read cloud approach to diverse environmental samples, especially for those in which isolation and culture have been limited in providing representative organisms, will help provide insight for the vast microbial life that is currently known. The genome drafts obtained could also be cross-referenced with available 16S rRNA amplicon sequencing data, to allow further characterization of microbiome samples for which taxonomic composition, but not functional characterization, is readily available.

Though comparative analysis on human genomes is now routine, comparative analysis of bacterial genomes and microbial metagenomes has lagged much further behind. The current reliance on incomplete reference sequence collections and highly fragmentary metagenome drafts obtained from conventional short read techniques is a core limitation in being able to perform such comparative analyses. Our work, which describes a shotgun metagenomic sequencing approach that successfully provides complete and high quality microbial genome drafts, does so at a price point that gives it relevance to the broader microbiome community. We anticipate that applications of read clouds to longitudinally collected samples will allow detailed comparative analyses. This will facilitate investigation of bacterial genomic plasticity through the identification of specific genomic alterations that individual bacteria acquire in response to selective pressures imparted on whole microbial communities. We anticipate that our approach will be a significant step forward in enabling comparative genomics for bacteria, enabling fine-grained inspection of microbial evolution within complex communities.

## Methods

### Healthy subject recruitment

Two healthy adult volunteers were recruited at Stanford University under an IRB-approved protocol (PI: Dr. Ami Bhatt). Informed consent was obtained. The subjects had no gastrointestinal disease or antibiotic use in the 6 months prior to sample collection.

### Sample Collection

#### Healthy volunteer stool samples

A single stool sample was obtained from each of the two healthy volunteers. Stool samples were placed at 4°C immediately upon collection, and processed for storage at −80°C the same day. Stool samples were aliquoted into 2mL cryovial tubes with no preservative. Samples were stored at −80°C until extraction.

#### Marine sediment sample

A deep-sea sediment core was collected using an MC-800 multicorer aboard the R/V Oceanus (expedition #1703A) 115 km off the coast near San Francisco, CA, USA in March of 2017 (36.61°N, 123.38°W; water depth 3535 m). The core was stored at 4C until extruded and sectioned within 24 hours of collection. Approximately 2g of sediment was sampled from the top 2.5 cm of sampled core using a cut-off syringe, flash frozen in liquid nitrogen, and stored at −80C until extraction.

### DNA preparation

#### Healthy volunteer stool samples and mock metagenome sample

DNA was extracted from Participant 1 (P1) stool with the Qiagen Gentra Puregene Yeast/Bacteria kit according to the manufacturer’s standard protocol with two modifications: a chilling step at −80°C for five minutes prior to DNA precipitation, and DNA precipitation with 14,000g, 20 minute centrifugation at 4°C. DNA was extracted from Participant 2 (P2) stool with the Qiagen QIamp Stool Mini Kit according to the manufacturer’s standard protocol, modified with an additional step after addition of buffer ASL. The additional step was 7 cycles of alternating 30 second periods of beating with zirconia beads in a Minibeadbeater (Biospec Products, Bartlesville, OK) and chilling on ice. DNA concentration was measured using Qubit fluorometric quantitation (see supplementary table 4 for measured concentrations).

DNA that was to be taken forward for to 10X Chromium preparation was size-selected with the BluePippin instrument targeting the 10kb-50kb size range, the maximum yielding measurable output. DNA for the SLR library preparation was size-selected with the BluePippin instrument targeting the 8-12kb size range as per the manufacturer’s recommended protocol. DNA for Truseq conventional short read library preparation was not size-selected. DNA from ATCC 20 Strain Staggered Mix Genomic Material was used directly without size selection for the mock metagenome. Libraries were prepared for sequencing with the 10X Genomics Chromium (10X Genomics, Pleasanton, CA), the Illumina Truseq SLR kit, or Illumina Truseq Nano kit according to the respective manufacturer’s standard protocol. Library fragment size was quantified with the Agilent 2100 Bioanalyzer instrument (Agilent Technologies, Santa Clara, CA) using the High Sensitivity DNA kit.

#### Marine sediment sample

DNA was extracted using the RNeasy PowerSoil DNA elution kit (Qiagen, Hilden, Germany; cat. no. 12867-25) in combination with the RNeasy PowerSoil Total RNA kit (Qiagen, Hilden, Germany; cat. no. 12866-25). The protocol was modified from the manufacturer’s instructions to include a bead-beating step of 5.5m/s for 2X 45s using a FastPrep-24 (MP Biomedicals, Santa Ana, CA, USA; cat. no. 116005500). DNA was eluted in 100l DNase, RNase-free water and stored at -80C until further processing. DNA was then size-selected with the BluePippin instrument targeting the 10kb-50kb size range (the maximum yielding measurable output), and a library was prepared for sequencing with the 10X Genomics Chromium (10X Genomics, Pleasanton, CA), according to the manufacturer’s standard protocol.

### Sequencing

#### Chromium libraries

Chromium libraries from the mock metagenome, healthy stool samples, and ocean sediment were sequenced with 2x151bp sequencing on an Illumina HiSeq 4000. The healthy stool samples were allocated a half lane each. The marine sediment was allocated a quarter lane and a full lane. The mock metagenome was allocated one lane. (See Supplementary Table 3 for total Gb coverage). Resulting sequences were demultiplexed and barcoded with the 10X Longranger mkfastq tool to generate raw reads, then subjected to quality control.

#### Truseq libraries

DNA from the healthy stool samples was prepared for sequencing with the Illumina Truseq library prep kit according to the manufacturer’s standard protocol and subjected to 2x101 bp sequencing on an Illumina HiSeq 4000. Each library was allocated a half lane of sequence coverage (see Supplementary Table 3 for total Gb coverage). Raw reads were then subjected to quality control (see below).

#### Synthetic long read libraries

DNA from the healthy stool samples was prepared for sequencing with the Illumina Truseq Synthetic Long Read library prep kit according to the manufacturer’s standard protocol. These libraries use the sample barcode to identify the 384 molecular partitions, so samples cannot be multiplexed. Thus, each library was necessarily allocated one full lane of 2x151 bp coverage on an Illumina HiSeq 4000 (see supplementary Table 3 for total Gb coverage). Raw reads were then subjected to quality control (see below).

### Quality control

Following sequencing, all libraries were trimmed using cutadapt^48^ using a minimum length of 60bp and minimum terminal base score of 20 (with the exception of the ATCC mock metagenome reads, which were trimmed with a minimum trimmed read length of 80bp and minimum terminal base score of 35, as well as 8bp removed from the 5’ end and 15bp removed from the 3’ end due to low read quality). Reads were synced and orphans (reads whose pair mates were filtered out) were placed in a separate single-ended fastq file with an in-house script.

### Assembly of mock metagenome and human stool samples

Data from Chromium and Truseq libraries were assembled using MetaSPAdes v3.11.1 ^49^ with default parameters. For Chromium libraries, MetaSPAdes assembled seed contigs were then assembled with Athena (see Supplementary Methods).

Synthetic long reads were assembled from trimmed sequencing reads with TruSPAdes^50^ with default parameters. Assembled synthetic long reads were then assembled into contigs with CANU ^51^ with the following parameters: errorRate=0.06, genomeSize=45.00m, contigFilter=“2 2000 1.0 1.0 2”, stopOnReadQuality=false.

### Assembly of marine sediment sample

Data from the marine sediment Chromium library was assembled using MEGAHIT^52^ with default parameters. MEGAHIT short-read assembled contigs were then used as seed contigs and assembled with Athena (see Supplementary Methods).

To make Athena assembly tractable on complex metagenomes, Athena was modified to only perform subassembly for well-covered seed contigs with a minimum short read sequence coverage of 20x. MEGAHIT contigs excluded from Athena assembly were then mapped back to the initial Athena draft, and each of these contigs was included in the final output if more than 2000 bases did not align to the initial draft.

### Assembly classification, genome draft binning, and gene identification

For each approach, raw short reads were aligned to assembled contigs with BWA^53^ to generate contig coverage profiles. Contigs were then binned with Metabat^18^ to form genome drafts. Bins were evaluated with Metaquast^54^ for assembly size and contiguity, CheckM^55^ for completeness and contamination as genome drafts, Prokka^56^ for gene content, Aragorn^57^ to count tRNA sequences, and Barrnap^58^ to count 5S, 16S and 23S ribosomal RNA loci. We adopt previously described standards defining a “medium quality” genome as one with at least 50% completeness and at most 10% contamination, and a “high quality” genome as one containing at least 18 tRNA loci, at least one copy each of 5S, 16S and 23S, at least 90% completeness and at most 5% contamination^47^.

Individual contigs from all assemblies were assigned taxonomic classifications using Kraken^59^ with a custom database constructed from the Refseq and Genbank^60, 61^ bacterial genome collections. Each genome draft was assigned a species-level label if >=60% of total bases within the draft shared a species-level classification. Otherwise, drafts were assigned the majority genus-level label.

### Code availability

The Athena assembler together with a demonstration dataset can be found at https://github.com/abishara/athena_meta. This example contains a subset of the read clouds from the ATCC 20 mock metagenome, for which assembly with Athena yields the full *Lactobacillus gasseri* genome in two sequence contigs.

### Data availability

The datasets generated during the current study are available in the NCBI Sequence Read Archive under Bioproject accession PRJNA380276. 10X read barcodes have been encoded as read groups for SRA archival, and must be reformatted as barcodes (read groups replaced with bar codes in individual reads) for use with Athena.

## Acknowledgements

The authors would like to thank Ekaterina Tkachenko for assistance preparing Truseq libraries, and Michael Snyder and members of the Bhatt lab for helpful feedback. The authors would also like to thank Hongxia Xu at Illumina for sharing read cloud sequencing data of ATCC 20 for the mock metagenome. This work was supported by NCI K08 CA184420, the Amy Strelzer Manasevit Award from the National Marrow Donor Program, and a Damon Runyon Clinical Investigator Award to A.S.B. E.L.M. was supported by National Science Foundation Graduate Research Fellowship DGE-114747. A.B. was supported by the Stanford Genome Training Program (SGTP; NIH/NHGRI) and the Training Grant of the Joint Initiative for Metrology in Biology (JIMB; NIST). A.E.D. was supported by National Science Foundation OCE-1634297. A.P. was supported by the Center for Dark Energy Biosphere Investigations Postdoctoral Fellowship. Access to shared compute resources was supported in part by NIH P30 CA124435 using the Stanford Cancer Institute Shared Resource Genetics Bioinformatics Service Center.

## Author contributions

A.B., E.L.M., A.S.B. and S.B. conceived of the study. Z.W. prepared read cloud libraries. E.L.M. extracted DNA and prepared SLR sequencing libraries. E.L.M. performed PCR validation, and Sanger sequencing. A.B. and S.B. conceived of the assembly approach. A.B. implemented the Athena assembler. M.K. modified the Flye assembler for use with Athena. A.P. and A.D. collected marine sediment samples and helped in analysis of these samples. A.B, A.S.B. and E.L.M. carried out all analyses, wrote the manuscript, and generated figures. All authors commented on the manuscript.

## Competing financial interests

The authors declare no competing financial interests.

## Supplementary Material

### Supplementary Discussion on Read cloud and SLR sequencing

Both synthetic long read (SLR) sequencing and read cloud sequencing leverage barcoding of short fragments derived from partitioned long input DNA fragments. The two approaches differ in how they trade off physical coverage of the target metagenome in long fragments (C_F) versus the coverage of each long fragment with short reads (C_R) given a fixed short read budget (C = C_F × C_R). Previous work using SLR sequencing to improve upon metagenomic assembly utilized high short-read coverage of individual long fragments (C_R = 50x). Illumina Truseq Synthetic Long reads are currently the only commercially available kit that enables SLR sequencing and it achieves partitioning with use of 384 well plate. The read cloud approach instead uses much lower short-read coverage of individual long fragments (C_R=0.1x) to allow each microbial genome to be covered with many more long fragments (Figure A.1). This is accomplished by using a much larger number of individual droplet partitions. The 10X Genomics platform, which we used to generate read clouds, do not perform within-partition amplification efficiently enough to allow high short-read coverage of individual long fragments for SLR sequencing.

The read cloud and SLR modes of operation necessitate different genome assembly approaches in order to achieve improved metagenomic drafts (shown in Figure A.1). Read cloud assembly with Athena *jointly* assembles short reads from multiple barcodes (see Methods section on Athena). Metagenomic assembly with SLR sequencing by contrast is a two stage process that first requires pre-assembly of raw barcoded short reads into virtual long reads for each partition; these long reads are then are then overlap layout consensus (OLC) assembled into much longer contigs.

**Figure A.1:**
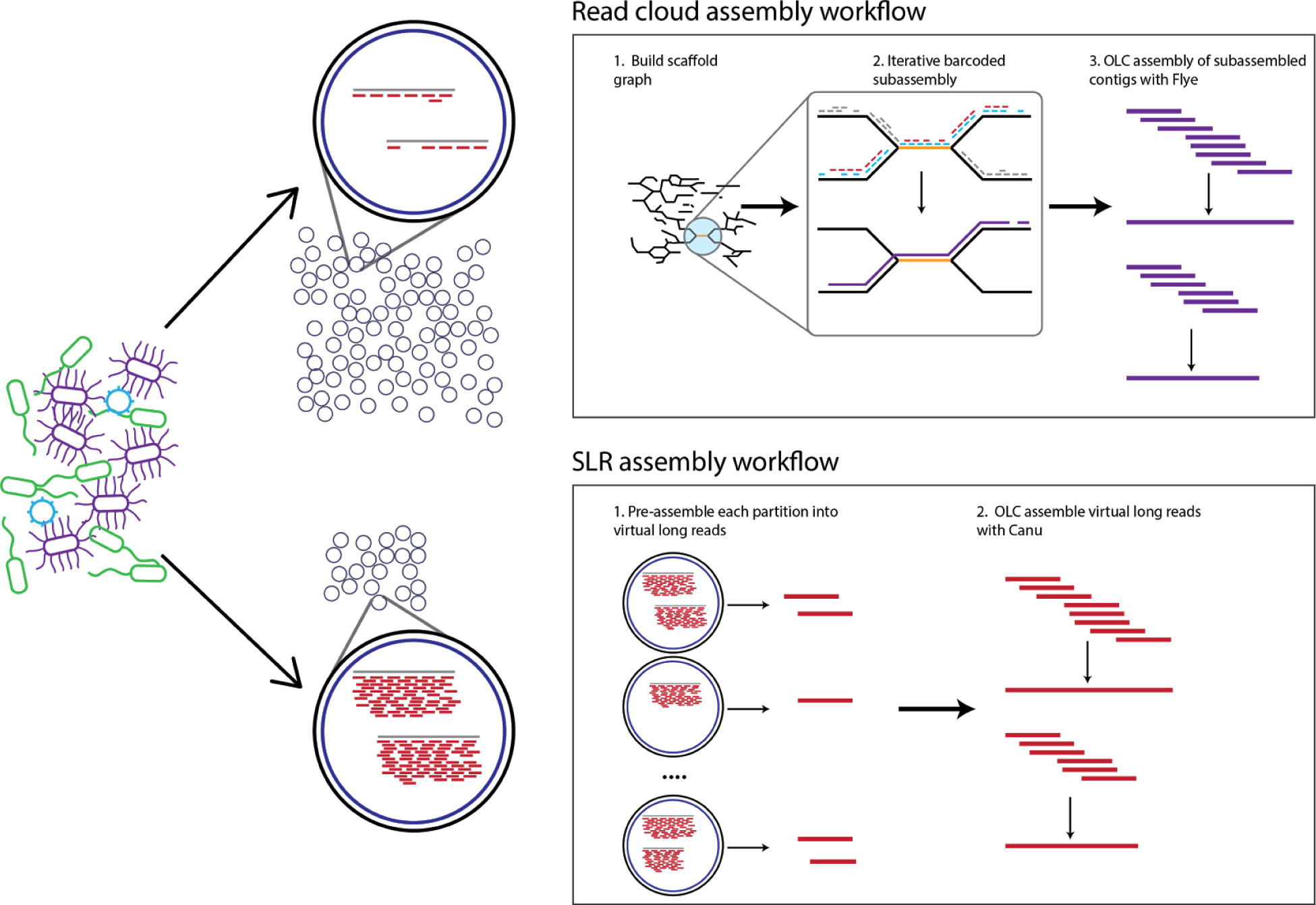
Read cloud and SLR sequencing and assembly workflows.

The read cloud approach allows assembly of more microbes than the SLR approach with a fixed total short-read sequencing budget C (Figure A.2). For a lower abundance microbe to be covered at C=50x, read clouds will sufficiently cover it with many long fragments (C_F=500x), each with shallow short-read coverage (C_R=0.1x). Athena read cloud assembly is able to jointly assemble these barcoded short reads to yield high quality drafts in this mode of operation. By contrast this same microbe covered at C=50x with the SLR approach, will have very low long fragment coverage (C_F=~1x) in the form of pre-assembled virtual long reads (C_R=50x). These long reads will not contain sufficient overlaps, and as a result the SLR assembly of this organism will be incomplete and contain many gaps. The read cloud approach allows assembly of many more community members in complex metagenomic samples over the SLR approach, effectively increasing the throughput 50-fold.

**Figure A.2:**
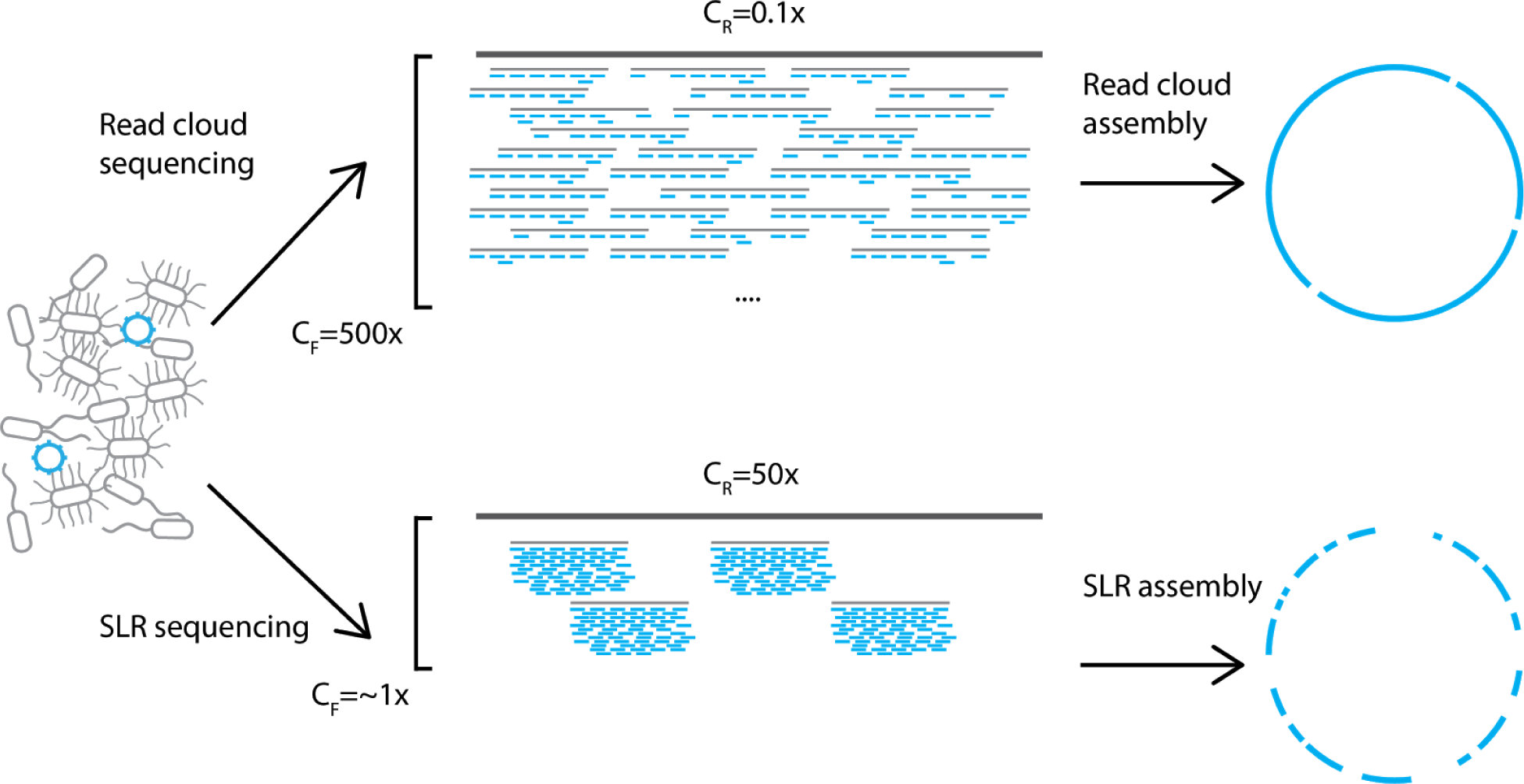
Overall throughput tradeoff between read clouds and SLRs under a fixed short-read sequencing budget C.

Several studies have utilized SLR sequencing for metagenomic sequence analysis. A previous effort to assemble a sample derived from the human gut, required 200Gb of raw short-read sequence to generate roughly 8Gb of virtual long read sequence. Further OLC assembly of these virtual long reads allowed assembly of some microbial community members present in the human gut into larger contigs^30^. Previous efforts have also been made to apply SLR sequencing to environmental microbiome samples, but these efforts all encountered difficulty in further assembling the virtual long reads into longer contigs^62, 63^. The lack of improvement in assembly draft quality is most likely due to higher species richness and evenness in representation in these samples, which leads to individual genomes being covered with few virtual long reads. The throughput afforded by our read cloud approach will facilitate assembly even in these highly diverse samples.

## Supplementary Figure and Table Legends

**Supp. Figure 1**

PCR and Sanger validation of several rRNA loci within the Athena assembly of two organisms within the ATCC mock metagenome, *A. baumannii* and *P. acnes*, found to be discordant with the reference assembly. a) PCR was performed with primers flanking rRNA loci as assembled in the Athena assembly (expected amplicon size: 7-8kb). Successful amplification was obtained, indicating misassembly in the existing reference sequence. PCR products were loaded on a 1% agarose gel alongside a 1kb+ DNA ladder. Lanes 2, 3, 4, and 5 contain bands for all four of the *A. baumannii* operons assembled by Athena. Lane 1 contains a band for one of the three *P. acnes* operons assembled by Athena; the others did not amplify. b) Sanger reads generated from each terminus of each of the five amplicons. Sanger reads crossing the 5’ and 3’ ends of the 16S/23S locus, as well as the location of the rRNA locus itself, are indicated by grey bars.

**Supp. Figure 2**

Genus-level community composition of DNA extracted with different conditions from the two healthy human gut samples P1 and P2. For each sample, DNA was extracted using either bead beating lysis or enzymatic lysis, and then subjected to a possible 10kb size selection as described in Methods. Sequencing libraries for DNA resulting from different preparations were then prepared, and the resulting short reads were classified to obtain community composition information. Genus-level community compositions appear concordant across different preparation conditions, with some enrichment of lower-abundance genera with enzymatic lysis and 10kb size selection.

**Supp. Figure 3**

Total assembled sequence within contigs of a minimum size for the three tested approaches for gut communities in samples P1 and P2. Only contigs with a minimum size of 100kbp are displayed. Read clouds placed substantially more sequence into larger contigs than compared to either SLRs or short reads. For sample P1, read clouds produced 64Mb of sequence within contigs of size >500kb, as compared to 11Mb from SLRs and 5Mb from short reads. For sample P2, read clouds produced 17Mb of sequence within contigs of size >500kb, as compared to 3Mb from SLRs and 0.5Mb from short reads.

**Supp. Figure 4**

Overview of analysis workflow performed on assembled sequence data. Reads are first assembled into metagenomic drafts. These sequence contigs are grouped into genome bins with Metabat^18^. Genome bins are then assessed as genome drafts under various quality criteria by their presence of single copy core genes as predicted by CheckM^55^. Genome bins are then taxonomically annotated at the species-level when possible and at the genus-level otherwise using contig classifications from Kraken^59^.

**Supp. Figure 5**

Kmer spectra (k = 31) derived from short reads in the read cloud library from sample P2, which were assigned to one of three organisms, *P. copri, Phascolarctobacterium* sp., and *S. wadsworthensis*, as described in Methods. Kmer counts were rarefied to equalize per-organism totals. *P. copri* possesses a subset of kmers occurring at a much higher incidence than the other two organisms assessed. Isolation those high abundance kmers from *P. copri*, and subsequent assembly of these kmers, yielded the sequences of high copy repeat elements present in this organism as described in Supplementary Methods.

**Supp. Figure 6**

Normalized histograms of long fragment lengths used in read cloud sequencing of P1 and P2 healthy participant guts. Fragments were estimated bioinformatically as described in Supplementary Methods. Long fragments of P1 were noticeably longer than those of P2, with median estimated long fragment lengths of 8161bp and 5071bp respectively.

**Supp. Figure 7**

Total assembled sequence within contigs of a minimum size for both Athena and conventional short-read assembly of the read cloud library of a marine sediment sample. Only contigs with a minimum size of 10kbp are displayed. As with the human gut samples, Athena assembly placed substantially more sequence into larger contigs than compared to conventional short-read assembly.

**Supp. File 1**

Genus-level short read classification of all samples. Kraken^59^ was used to classify short reads from all samples sequenced on the Illumina HiSeq 4000 (see Methods). Figure 2a,b and Supplementary Figure 5 are based on these classifications. All taxonomic classifications occurring in at least 1% of reads are presented. Column format is as follows:

1. Participant
2. Library
3. Number of reads classified within this taxon (and all child taxa)
4. Number of reads classified specifically to this taxon (occurs when a more specific classification is not possible)
5. Classified taxon

**Supp. File 2**

Taxonomic classifications of contigs assembled by the three approaches from the two healthy human guts obtained with Kraken^59^.

**Supp. File 4**

Assembled metagenome drafts for the two healthy human gut samples P1 and P2 across three sequencing and assembly methods.

**Supp. File 5**

Taxonomic classifications of contigs from both short-read assembly and Athena assembly obtained with Kraken59 for the marine sediment sample.

**Supp. File 6**

Assembled metagenome drafts for the marine sediment sample obtained from using conventional short-read assembly and Athena assembly. Contigs less than 1kb in size are excluded.

**Supp File 7**

Assignment of metagenome draft contigs to individual bins obtained from each of the three approaches applied to the two healthy human gut samples.

**Supp. Table 1**

Per-species MetaQUAST statistics for Athena read cloud assembly and standard short-read assembly run on ATCC 20 staggered DNA mixture (mock metagenome).

**Supp. Table 2**

Per-species counts of 16S and 23S rRNA units in ATCC 20 closed genome reference sequences, and counts of assembled 16S and 23S instances using Athena read cloud and standard short-read assembly on ATCC 20 staggered DNA mix. Instances of 16S and 23S units appearing within 2000kbp of each other are counted as a full rRNA operon, otherwise these units are counted separately. Only units with at least 3kbp of flanking sequence assembled on both sides are reported.

**Supp. Table 3**

Concentration and total mass of DNA samples before and after size selection.

**Supp. Table 4**

Total reads and sequencing coverage for all libraries before and after quality control.

**Supp. Table 5**

Assembly statistics, completeness metrics, tRNA and rRNA loci counts and total annotated genes from all sequencing and assembly approaches for all annotated species in the two healthy guts. Results are shown for the largest bin of each species and each is displayed only if at least one approach obtained a complete bin.

**Supp. Table 6**

Sizes, estimated copy numbers, and annotations of high copy repeat elements discovered in *Prevotella copri* in sample P2.

**Supp. Table 7**

Sequence identity and percent coverage for chosen references for 17 organisms that were assembled well with read clouds. In addition, the N50 values for the reference and read cloud assemblies, as well as the reference quality (closed or incomplete) are given.

**Supp. Table 8**

Mapping between contigs and bins for healthy gut assemblies.

**Supp. Table 9**

Mapping between contigs and bins for ocean sediment assemblies.

## Supplementary Athena Description

### Overview

We developed Athena to perform metagenomic assembly using barcoded short-read sequences (100bp) derived from partitioned long input DNA fragments (5-20kbp), which we refer to as read clouds. In this study, we apply Athena to read cloud datasets generated with the 10X Genomics Chromium instrument. In principle, the long fragments that are used as input to these platforms allow resolution of repeats contained within these fragments. The length of long fragments used exceeds the length of a typical bacterial repeat (rRNA operon of size ~5kbp). However, the barcode-specific coverage of each long fragment is too sparse to allow *de novo* assembly of each in isolation. Furthermore, the long range information encoded within the raw output of each barcode, which is in the form of unordered and unoriented short-read sequences, does not fit well into existing sequence assembly algorithms.

Athena uses the barcode information to perform metagenomic assembly by proposing a series of simplified assembly tasks, each of which can be performed using existing assemblers as black box subroutines. In this section, we introduce the concept of barcoded subassembly, a new approach to utilize read clouds for short-read sequence assembly. We then provide an overview of the following steps Athena performs for metagenomic assembly:

1. *Initial assembly graph construction:* First an existing short-read assembler (metaSPAdes) is applied to the input short reads stripped of the barcodes to obtain an initial sequence covering of the underlying metagenome in the form of (possibly short) sequence contigs. A scaffold graph is then constructed using the paired-end information from short-read alignments to these contigs. This scaffold graph contains branches that can be attributed to nearly identical repeats, small divergent sequences between otherwise identical strains, or conserved sequences.
2. *Iterative barcoded subassembly:* Barcoded short reads are then mapped to this scaffold graph. This scaffold graph is then traversed to propose *many* smaller input read subsets, each to be used in a much simpler assembly problem (barcoded subassembly). The resulting subassembled contigs yield unambiguous paths through the scaffold graph.
3. *Overlap assembly of subassembled contigs:* The resulting subassembled contigs are then passed as reads to the long read de Bruijn graph based assembler, Flye^39, 40^, for further assembly of these much larger overlapping sequences.

### Barcoded subassembly overview

In order to use the barcodes to identify true paths through complex whole genome assembly graphs, we adopted an approach to pool sub-selected groups of barcoded reads to create smaller local assembly problems, each simple enough to be solved using off-the-shelf short-read assemblers. We define *barcoded subassembly* to be the pooling of barcodes that locally cover a desired genomic seed region to further assemble sequence in this region.

We provide the following example to motivate this approach. Consider a read cloud experiment of a whole genome with long input DNA fragment sizes of <25kb, long fragment coverage of C_F = 500x, and barcode short-read coverage of C_R = 0.1x. For any particular small seed region of the genome, there are roughly 500 barcodes containing long fragments -- and therefore short reads -- that cover this region. Though each of these barcodes in isolation provides only sparse coverage, pooling of all short reads in these barcodes sufficiently covers a local seed sequence at 0.1 × 500 = 50x. Furthermore, given the long input fragments, coverage of flanking sequence near the seed will also be 50x tapering off to zero as distance from the seed increases to 25kb. The pooling of short reads from these subassembly barcodes allows isolation of this local target sequence from the broader whole genome assembly problem. It also allows us to leverage well-established short-read assemblers as a black box subroutine to reconstruct this target sequence de novo. Figure B.1 provides an overview of barcoded subassembly.

**Figure B.1:**
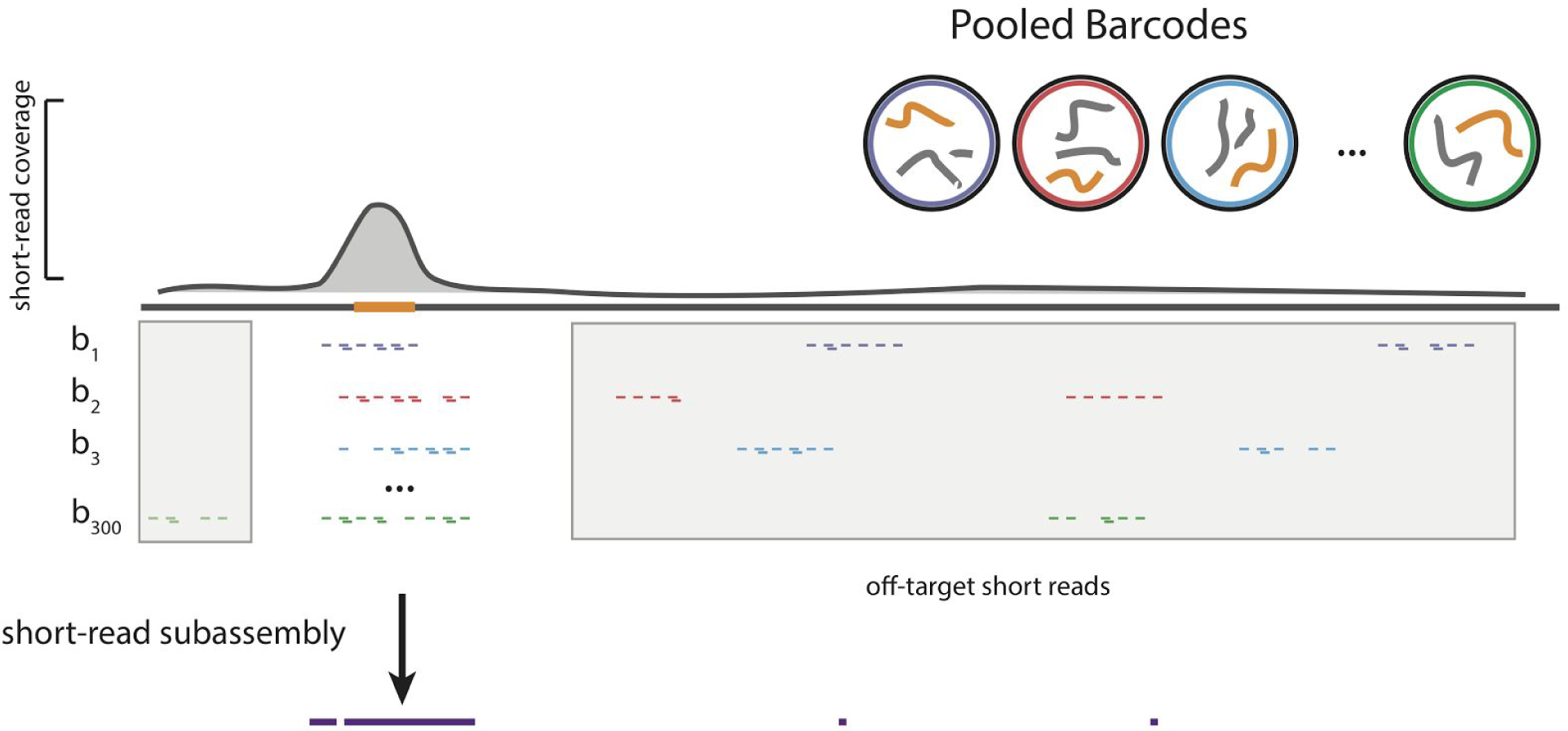
Example subassembly of a local target sequence from many sparsely covered barcodes.

The long fragment partitioning in current read cloud platforms is imperfect, i.e. there is often more than one fragment in each partition. The most recently developed 10X Genomics platform, which leverages the highest number of partitions to date, still has an average of 10 long fragments present within each partition. Consequently, roughly 90% of all short reads pooled in a subassembly will actually originate from *off-target* sequence of the whole metagenome (as shown in Figure B.1). Off-target reads are randomly distributed throughout the whole metagenome such that any sequence other than the target should be covered at <1x and remain unassembled.

### Athena: Initial assembly graph construction

Athena first uses the short-read assembler metaSPAdes^64^ to obtain an initial sequence covering of the underlying metagenome in the form of (possibly short) sequence contigs, which we refer to as seeds. Raw barcoded reads are first stripped of their barcode information and passed to metaSPAdes as is for short-read assembly. The raw reads are mapped back to these seed contigs, and paired end mappings that span two seed contigs are considered for edge creation in a scaffold graph. The 10X Genomics platforms have a higher chimeric read fragment rate, such that a significant fraction of read pairs map with an intervening distance that exceeds the expected library fragment size. In order to prevent these from introducing spurious connections in the scaffold graph, we perform the following steps:

1. For any two seed contigs that are still connected by at least three spanning read pairs, the mapped positions of these spanning read pairs on each seed contig are clustered together into 500bp neighborhoods, corresponding to the average library fragment size.
2. All clusters are examined and if any single cluster on each seed contig contains more than 50% of these spanning read pairs, then an edge is added in the scaffold graph. Otherwise, the candidate edge is assumed to be spurious and discarded. This filtering process greatly reduces the number of proposed subassemblies to perform.

This scaffold graph contains branches that can be attributed to nearly identical repeats, small divergent sequences between otherwise identical strains, or largely conserved sequences. Raw reads annotated with the barcode information are then mapped back to this scaffold graph in order to propose read subsets for subassembly.

### Athena: Iterative barcoded subassembly

Athena uses the barcode information to propose a series of simple subassembly tasks that can be performed using existing short-read assemblers as black box subroutines. For every edge between two seeds in the constructed scaffold graph, subassembly of these linked seed contigs is performed with the following steps:

1. Barcodes containing at least one read mapping to both seeds within 10kb of the scaffold edge connection point are selected as candidates for subassembly. Pooled reads from these candidate barcodes potentially contain contiguous sequences that bridge together the two seed contigs.
2. Pooled reads that map to these two seeds are used to estimate short-read coverage of the target sequence within the subassembly. If the short-read coverage is estimated to be low (<10x), then this subassembly is skipped as the local target is unlikely to assemble at low depths. If the short-read coverage is estimated to be high (>100x), then the subassembly barcodes are first downsampled to accelerate subassembly and minimize unnecessary pooling of off-target reads.
3. If downsampling is required, barcodes are sorted in decreasing order of number of reads mapping to these two desired sequence contigs. Barcodes are added for subassembly until the estimated local coverage of the two seed contigs exceeds 100x. Preferential inclusion of barcodes with the highest number of reads mapping to the seeds corresponds to selection of the longest source fragments covering this region.
4. All raw reads from the selected subassembled barcodes are then pooled together and assembled with a modified version of IDBA-UD^65^, IDBA-SUBASM (described below), to yield subassembled contigs.

Figure B.2 shows that inclusion of all barcodes with at least one read mapping to a single seed (shown on the right in orange) in the scaffold graph, may not help to separate short reads originating from different microbial strains. This single seed sequence may be shared across multiple microbial strains such that inclusion of all barcodes would not help identify a single unique path. Subassembly of pooled reads from all barcodes would likely yield contigs that break precisely at the locations of the scaffold connections. However, when the criterion for subassembly barcodes is changed to allow only barcodes with at least one read mapping to both a seed and also a single neighbor seed contig in the scaffold graph, then the pooled short-reads can carve out an unambiguous path (as shown in Figure B.3). Provided these pooled barcodes have high enough coverage, short-read assembly of the short-reads would instead yield contigs that assemble through branches in the scaffold graph. The resulting subassembled contigs are likely to disambiguate other branches in the scaffold graph because the pooling of *all* reads within the chosen barcodes also draws in reads from flanking regions due to the long input DNA fragments.

**Figure B.2:**
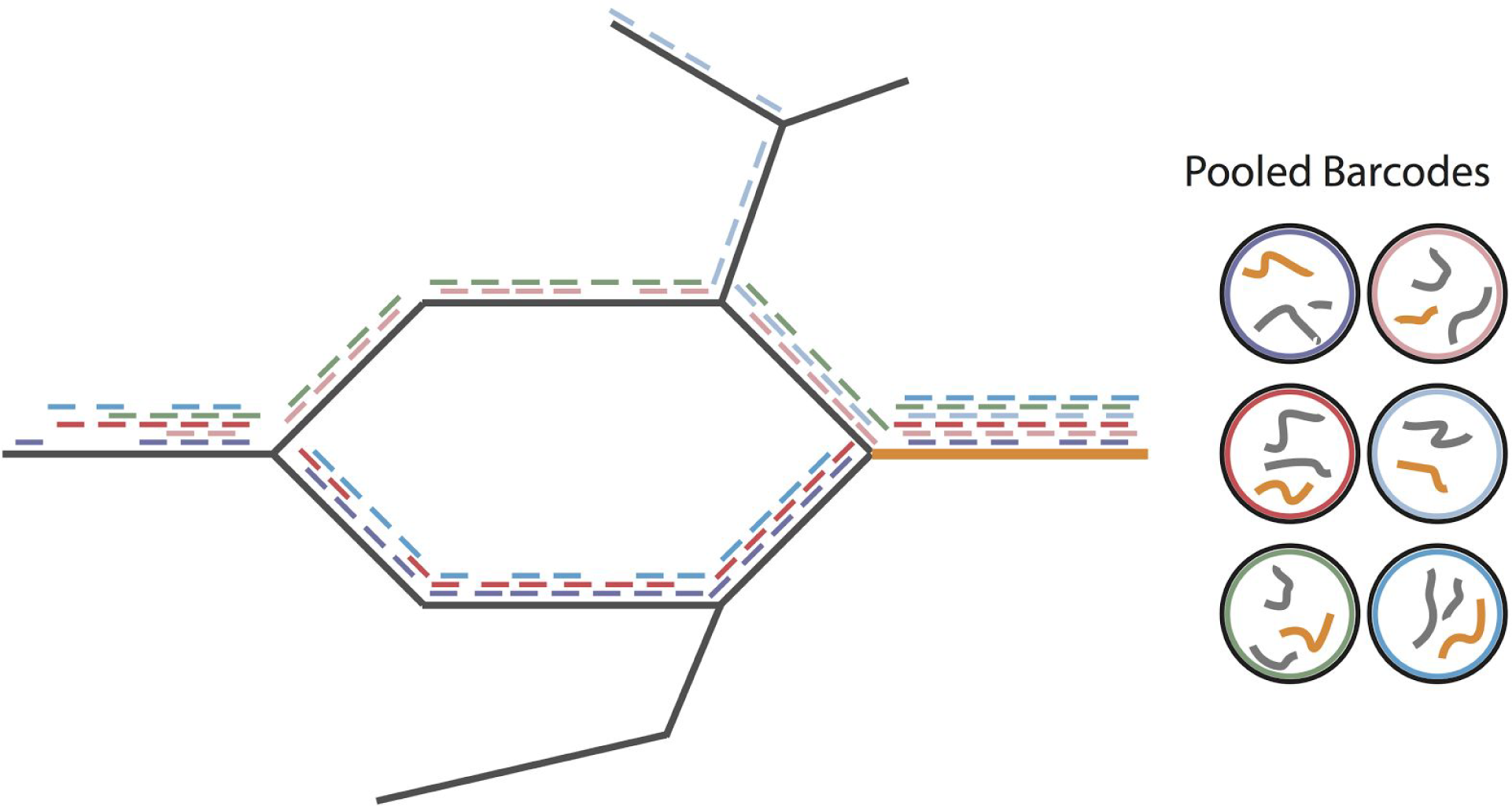
Example of an uninformative choice of subassembly barcodes

**Figure B.3:**
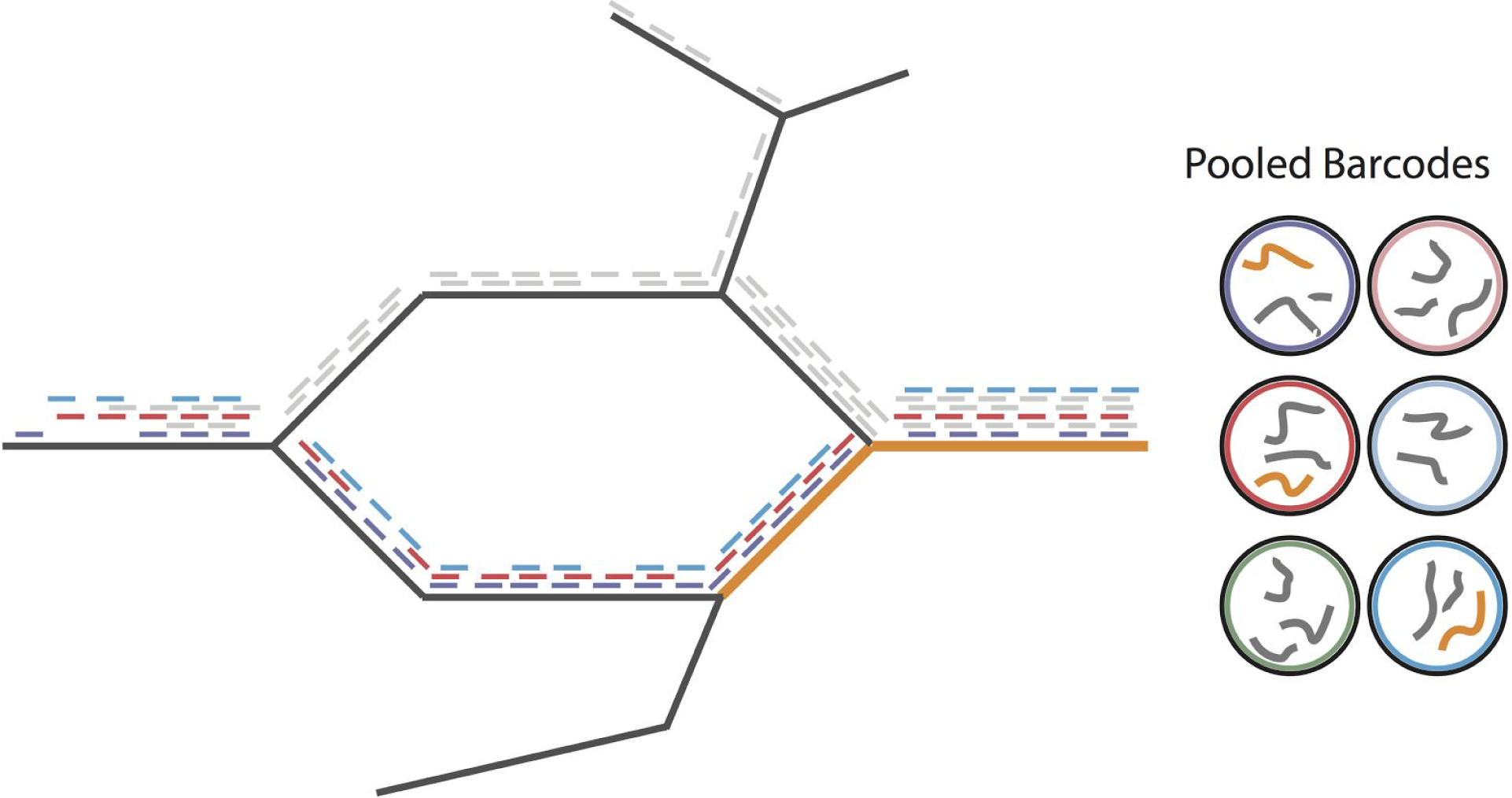
Example of a well chosen subassembly barcodes carving an unambiguous path

In addition to subassemblies induced by edges in the scaffold graph, each seed contig can give rise to up to two additional subassemblies. If a seed is small (<= 20kb), then barcodes with at least one read mapping to it (downsampled as described above), are pooled for subassembly. If a seed is large (>20kb), then two subassemblies for it are performed: one for the head 10kb and one for the tail 10kb. Intuitively, we only seek to extend the ends of large sequence contigs as the middle sequence has largely been resolved by short-read assembly alone. Large seeds contain a very large number of barcodes with at least one read mapping to them, and if pooled together may bring in too many off-target reads to allow for accurate reconstructions of the ends.

Though pooling of subassembly barcodes draws in short-reads that sufficiently cover the local target sequence and its flanks, short-reads from off-target long fragments are also drawn in (as shown in Figure B.1). Barcodes are pooled until the local sequence is estimated to be covered at approximately 100x. The long fragment dilution and partitioning within the 10X Genomics platform is imperfect and each barcode contains on average roughly 10 long fragments, only one of which will be of interest when its chosen for subassembly. This implies that the majority (>90%) of reads will originate from off-target fragments. However, the “background” metagenome usually contains hundreds of different bacterial species in diverse samples such that these off-target fragments have a low probability of collision. Recall that although bacterial genomes have low repeat content in contrast to mammalian genomes, all bacteria share the conserved rRNA operon. This repeat unit is usually ~5kbp in size and each bacteria can have up to 20 identical copies of this repeat.

A quick back-of-the-envelope calculation reveals that pooling of subassembly barcodes to a depth of 100x almost surely pools enough reads from the rRNA operon such that spurious reconstructions of this repeat unit (and other ones of similar copy number profile) are possible. Each 10X barcode covers the long fragments at a depth of 0.1x, such that roughly 1000 barcodes need to be pooled to receive a local coverage of 100x. For our metagenomic samples, each 10X barcode contains on average 10 long fragments and produces on average 100 short-reads covering these fragments. As nine out of ten 10 fragments in the 1000 subassembly barcodes will be off-target, roughly 90% × 100 reads per barcode × 1000 barcodes = 90,000 short reads will be off-target. Assuming a 5Mb genome with 20 rRNA repeats of size 5kb, roughly 2% of all bacterial genomic sequence is rRNA operons. This implies that in every subassembly there is 2% × 90,000 = 1,800 of short-read rRNA sequence, yielding 1,800 × 150bp reads / 5kbp = 54x coverage of the rRNA operon.

Empirically we found these spurious repeat reconstructions to always be entirely disconnected from the De Bruijn Assembly Graph constructed in subassembly. We leveraged IDBA-UD, a short-read assembler designed for use with highly uneven short-read coverages, and added a new module IDBA-SUBASM to prune these reconstructions. Pooled short reads are first jointly assembled using the baseline IDBA-UD algorithm. The selected input seed sequences that gave rise to the subassembly are then aligned to the resulting De Bruijn Graph with kmer size 100. Any contigs arising from connected components that do not have at least 100bp of seed sequence alignment hits are then pruned before branch points in the graph are broken and contigs are produced. We found this simple procedure pruned nearly all spurious off-target reconstructions.

### Athena: Overlap assembly of subassembled contigs

The subassembled contigs, which contain large flanking overlaps (with unique as well as repetitive sequence), together with the initial seed contigs, are passed as reads to the long read assembler Flye^39, 40^. First, Flye constructs a draft assembly by greedily extending the sub-assembled contigs using the overlaps between them, without attempting to resolve repeats accurately. The draft assembly is then combined into the assembly (repeat) graph, in which repeats longer than 1000bp are revealed. Flye then maps the subassembled contigs back onto the assembly graph and simplifies it by resolving the repeats that are spanned in full by the subassembled contigs. The edges of the assembly graph are then output as final assembled sequences. The parameters of the original algorithm, which were originally designed for assembly of long and noisy reads, were modified to reflect the different error pattern and coverage distribution of subassembled contigs.

The resulting draft metagenome assembly contains more complete sequence contigs that do not have gaps and resolve shared sequences too difficult to assemble from short-read techniques alone (Figure B.4). Repeats that cannot be unambiguously spanned by subassembled contigs remain unresolved by the overlap assembler.

**Figure B.4:**
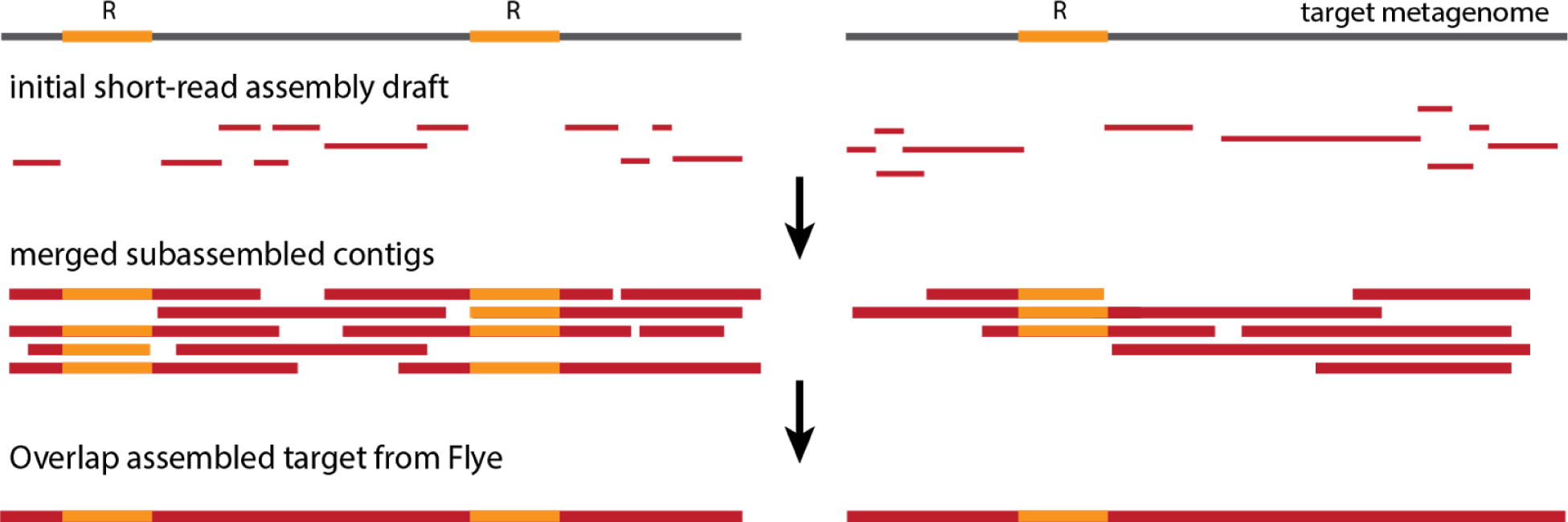
Flye overlap assembly of subassembled contigs.

## Supplementary Methods

### Prevotella enriched *k*-mer assembly

In order to ascertain the reason for poor assembly of *Prevotella copri* in all three trialled sequencing and assembly approaches, we performed a close examination of *k*-mers present in short reads originating from *P. copri* and two other organisms, *Phascolarctobacterium* sp. CAG:207 and *Sutterella wadsworthensis*, which were both well-assembled for comparison.

We first attempted to determine the set of short reads originating from each of these organisms, inclusive of those from potential repeat sequences that were not represented in the initial seed assembly draft. Subassembled contigs were used to for recruit short reads for each organism in order to account for repeats that may have been locally assembled by Athena (in the intermediate subassembled contigs) yet unresolved in the final output. Subassembled contigs were first assigned to each organism by mapping with BWA against all genome bins. Short reads were then mapped with BWA against all subassembled contigs, and assigned to the respective organisms on the basis of the subassembled contig assignments.

K-mer counts for each organism were generated from the read sets described above using Jellyfish^66^ with k = 31. *P. copri, Phascolarctobacterium* sp., and *S. wadsworthensis* were determined to have median coverages of 2836x, 48x, and 74x respectively. In order to determine the subset of k-mers originating from potential high copy repeats, we isolated overrepresented k-mers with occurrences exceeding 10 times the median coverage of that organism. These kmers were assembled using SPAdes^49^ with default parameters. *P. copri* was the only organism for which SPAdes yielded assembled sequence. For each predicted repeat, the multiplicity was determined by normalizing median repeat k-mer abundance by median *P. copri* coverage. This analysis yielded 1866bp, 1824bp, 1449bp, 631bp, and 507bp sequences with multiplicities 15, 23, 63, 13, and 13 respectively. These candidate repeats were next annotated using both nucleotide alignments with BLAST and translational alignments with BLASTx^67^. Two of the five repeats were annotated at the nucleotide level to known transposase sequences. However, all five repeats were annotated at the amino acid level to known transposase sequences (Supplementary Table 3.5).

### Read cloud fragment size estimation

In order to determine the effect of DNA extraction approach on read cloud size in finished 10X Chromium libraries, long fragment lengths were estimated from the BWA^53^ alignments of input short reads to assembled seed contigs. Only alignments to large input seed contigs (minimum size 50kbp) were used to avoid edge effects. For each barcode, short-reads were clustered as belonging to the origin same long fragment if the gap between alignments was less than or equal to 20kbp. The difference between the maximum and minimum alignment positions were used as the estimated long fragment size for each barcode cluster. Estimated fragment sizes for read cloud libraries from the two healthy human gut samples are shown in Supplemental Figure 2.

